# Theory on the looping mediated directional-dependent propulsion of transcription factors along DNA

**DOI:** 10.1101/418947

**Authors:** R. Murugan

**Affiliations:** Department of Biotechnology, Indian Institute of Technology Madras, Chennai, India.

**Keywords:** DNA-protein interactions, combinatorial binding, transcription factor binding, DNA-looping, *cis*-regulatory modules

## Abstract

We show that the looping mediated transcription activation by the combinatorial transcription factors (TFs) can be achieved via directional-dependent propulsion, tethered sliding and tethered binding-sliding-unbinding modes. In the propulsion mode, the first arrived TF at the *cis*-regulatory motifs (CRMs) further recruits other TFs via protein-protein interactions. Such TFs complex has two different types of DNA binding domains (DBDs) viz. DBD1 which forms tight site-specific complex with CRMs via hydrogen bonding network and the promoter specific DBD2s which form nonspecific interactions around CRMs. When the sum of these specific and cumulative nonspecific interactions is sufficient, then the flanking DNA of CRMs will be bent into a circle over the TFs complex. The number of TFs involved in the combinatorial regulation plays critical role here. When the site-specific interactions and the cumulative nonspecific interactions are strong enough to resist the dissociation, then the sliding of DBD2s well within the Onsager radius associated with the DBD2s-DNA interface towards the promoter is the only possible way to release the elastic stress of the bent DNA. The DBD2s form tight synaptosome complex upon finding the promoter via sliding. When the number of TFs is not enough to bend the DNA in to a circle, then tethered sliding or tethered binding-sliding-unbinding modes are the possibilities. In tethered sliding, the CRMs-TFs complex forms nonspecific contacts with DNA via dynamic loops and then slide along DNA towards promoter without dissociation. In tethered binding-sliding-unbinding, the CRMs-TFs performs several cycles of nonspecific binding-sliding-unbinding before finding the promoter. Elastic and entropic energy barriers associated with the looping of DNA shape up the distribution of distances between CRMs and promoters. The combinatorial regulation of TFs in eukaryotes has evolved to overcome the looping energy barrier.

## INTRODUCTION

Site-specific binding of transcription factors (**TFs**) at their *cis*-regulatory motifs (**CRMs**) on the genomic DNA in the presence of enormous amount of nonspecific binding sites is essential for the activation and regulation of several genes across prokaryotes to eukaryotes (1–3). Binding of TFs with their CRMs was initially thought as a single-step three-dimensional (**3D**) diffusion-controlled collision process. Kinetic experiments on *lac*-repressor-Operator system revealed a bimolecular rate in the order of ~10^9^-10^10^ M^−1^s^−1^ that is ~10-10^2^ times faster than the Smoluchowski type 3D diffusion-controlled rate limit (4). Berg et.al. (5, 6) successfully explained this inconsistency using a two-step mechanism by establishing the key concept that TFs first bind with DNA in a nonspecific manner via 3D diffusion and then search for their cognate sites via various one-dimensional (**1D**) facilitating processes such as sliding, hopping and intersegmental transfers. Here 1D diffusion with unit base-pair step-size of TFs is the sliding, few base-pairs (bp, 1 bp = *ld* ~ 3.4 × 10^−10^ m) step-size is called hopping and few hundred to thousand bps step-size is called intersegmental-transfer. Intersegmental transfers occur whenever two distal segments of the same DNA polymer come in nearby over 3D space via ring closure events (7–9).

Specific binding of TFs with DNA is affected by several factors (9) viz. **a**) conformational state of DNA (9, 10) **b**) spatial organization of various functionally related combinatorial CRMs along the genomic DNA (11, 12), **c**) presence of similar or other dynamic roadblock proteins (13) and semi-stationary roadblocks such as nucleosomes in eukaryotes (14–17), **d**) naturally occurring sequence mediated kinetic traps on DNA (18, 19), **e**) conformational fluctuations in the DNA binding domains of TFs (20–22) and **f**) the nonspecific electrostatic attractive forces and the counteracting shielding effects of other solvent ions and water molecules acting at the DNA-protein interface (23). Several theoretical models (8, 9, 18, 21, 24), computational (25–28) and experimental studies (29) have been carried out to understand the effects of factors **a-f** on the kinetics of site-specific DNA-protein interactions.

In general, the searching efficiency of TFs depends on the relative amount of times spent by them on the 3D and 1D diffusions (11, 21). Clearly, neither pure 1D nor 3D diffusion is an efficient mode of searching (9, 11). Under ideal situation, maximum searching efficiency can be achieved only when TFs spend equal amount of times in both 1D and 3D diffusions (8, 11). This trade off balance between the times spent on different modes of diffusions will be modulated by the factors **a-f**. For example, presence of nucleosome roadblocks warrants more dissociations and 3D excursions of TFs rather than 1D sliding (14). Sequence specific fast conformational switching of DNA binding domains between stationary and mobile states helps TFs to overcome the sequence traps (14). Relaxed conformational state of DNA enhances more sliding rather than hopping and intersegmental transfers and so on (9). Conformational dynamics of DNA also modulates the speed of gene activation and regulation. In this context, looping of DNA is critical for the activation and expression of various genes across prokaryotes to eukaryotes (3, 30-34). Combinatorial binding of TFs with their specific CRMs on the genomic DNA activates the downstream promoters of genes via looping of the intervening DNA segment to form a synaptosome type complex (1, 35). In most of the molecular biological processes, DNA-loops are warranted for the precise protein-protein interactions which are the prerequisites for transcription and recombination (36).

The statistical mechanics of looping and cyclization of linear DNA has been studied extensively in the literature (33, 37, 38). However, it is still not clear why DNA-loops have evolved as an integral part of the activation and repression of transcription and recombination although such underlying site-specific protein-protein and protein-DNA interactions can also be catered straightforwardly via a combination of 1D and 3D diffusions of TFs (5, 6, 21, 39). That is to say, upon arrival at the CRMs, TFs can directly slide or hop along the DNA polymer to reach the promoters. Schleif (31) had argued that the looping of DNA can simplify the evolution of the genomic architecture of eukaryotes by not imposing strict conditions on the spacing between the TF binding sites and the promoters. This is logical since a given set of TFs need to regulate several different genes across the genome. Therefore, placement of TF binding sites near a specific gene can be a disadvantage for other genes along the genomic evolution. Similarly, placement of TF binding sites near every gene is not an efficient genome design. The DNA loops also play critical roles in the transcription bursting (40) and memory (41). It is still not clear about **a**) the purpose of DNA loops in the transcription activation and **b**) the exact mechanism by which the DNA-loop is formed between the CRMs and promoters via TFs though Rippe et.al., (32) had already taken several snapshots of the looping intermediates. In this paper, we will show that the DNA-loops can propel TFs towards the promoters. Using computational tools, we further demonstrate that the looping mediated propulsion of TFs along DNA can actually help in finding the direction of the promoter region and also shape up the genomic architecture.

## THEORETICAL METHODS

### DNA loops mediated transcription activation in eukaryotes

Let us first list out the basic facts on the mechanism of distal action of CRMs-TFs system on the downstream promoters in the process of transcription activation.

a. Both theoretical investigations (5, 6, 8, 9, 21) and experimental observations (42, 43) suggested that TFs recognize their CRMs via a combination of 1D and 3D diffusions. The key idea here is that TFs scan a random piece of DNA via 1D diffusion after each of the 3D diffusion mediated nonspecific collisions (5, 9, 21). In contrast, the reacting molecules dissociate immediately upon each of their unfruitful collisions in the standard Smoluchowski model. When the dynamics of TFs is confined well within the *Onsager radius* of the DNA-protein interface, then it is categorized as the 1D diffusion. When TFs escape out of the Onsager radius and perform free 3D excursions, then we classify it as the 3D diffusion (9). The ***Onsager radius*** connected with the DNA-protein interface is defined (9) as the distance between the positively charged DNA binding domains of TFs and the negatively charged phosphate backbone of DNA at which the overall electrostatic interaction energy is same as that of the background thermal energy (equals to ~1 *kBT*) (Section 1, Supporting Materials). Further, one can define the dissociation of site-specific CRMs-TFs complex as the thermally induced separation of TFs from CRMs out of the Onsager radius associated with their interface.
b. The first arrived one of the combinatorial TFs further recruits other TFs (3, 44) around CRMs mainly via the cooperative protein-protein interaction among TFs and nonspecific electrostatic interactions between the DBDs of other TFs and DNA. This results in the formation of the CRMs-TFs complex. This is the well-established recruitment model (3) on the regulatory mechanism of the combinatorial TFs networks (45).
c. Transcription activation is achieved upon the distal communication between the CRMs-TFs complex with the corresponding promoter (1–3).
d. Binding of these combinatorial TFs at CRMs locally bends the DNA. At the end of these processes, synaptosome type DNA-loops connecting CRMs-TFs complex with the promoters are observed in most of the transcriptionally active genes of eukaryotes (3, 44).

Clearly, TFs activate transcription via two sequential steps viz. they bind their CRMs in the first step and then distally communicate with the promoter in the second step. To understand the role of DNA-loops, we consider two possible scenarios viz. looping mediated versus a hypothetical pure 3D1D diffusion mediated distal communication between CRMs-TFs and promoters. In both these scenarios, TFs locate their respective CRMs via a combination of 1D and 3D diffusion in the first step. They differ only in the second step where TFs dissociate from their CRMs and communicate with the promoters via a combination of 1D and 3D diffusions in the second case and, the distal communication will be through DNA-loops in the first case. We denote the search time required by TFs to locate their CRMs in the first step of transcription activation as *τ*_*S*_. Clearly, those factors **a-f** listed in the introduction section significantly modulate this quantity. We will not recalculate this here since enormous amount of literature already exists (Section 1 of the Supporting Materials) on the derivation of this quantity under various conditions (9, 18, 21, 46-48). In the following sections, we will compute the mean time required by the CRMs-TFs complex to communicate with the promoter via DNA-loops.

### Preliminary assumptions

Upon observing the open synaptic complexes of the transcriptionally active genes of eukaryotes with DNA loops, one can conclude that the combinatorial TFs complex which activates the transcription via DNA-loop has at least two different types DNA binding domains (DBDs) viz. DBD1 that corresponds to CRMs (Fig. 1) and DBD2 that corresponds to the promoter region. These two different DBDs may belong to two different TFs of the combinatorial TF complex. For example, in Fig 1, the DBD of TF-1 (DBD1 type) specifically binds CRMs and DBD of TF-4 (DBD2 type) specifically binds the promoter. Clearly, DBD1 type domains specifically interact with CRMs via hydrogen bonding network. Instead, the DBD2s of the other combinatorial TFs bind near CRMs region via non-specific electrostatic interactions and they are further stabilized by the protein-protein interaction network. For example, the tetrameric Lac I complex binds two different Operator regions that induces looping of DNA (2, 3). Further, the tetramers of repressor molecules bound at these two different binding sites communicate via protein-protein interactions among them. The DNA-loop is stabilized by an octamer form of the Lac I repressor protein. Such mechanisms are common in case of multiprotein mediated DNA-looping and transcription activation in eukaryotes. We further assume that TFs reach their specific binding site in the first step via a combination of 3D and 1D diffusions (5, 6, 8, 21, 49) in line with two-step site-specific DNA-protein interaction models and subsequently bends the DNA upon binding their specific sites located at the corresponding CRMs (32, 33).

**FIGURE 1.**
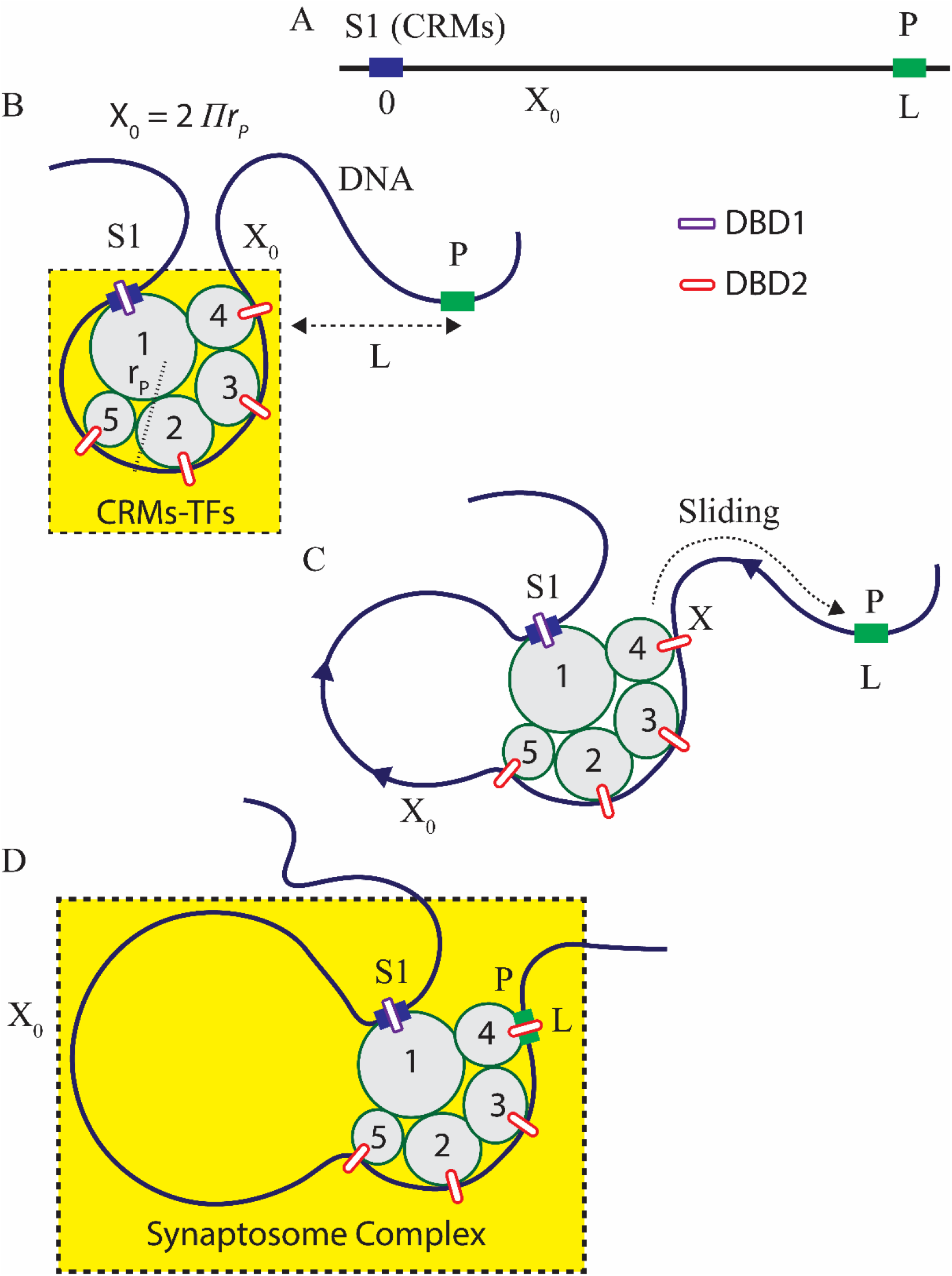
Directional-dependent propulsion of combinatorial TFs in the looping mediated transcription activation. **A**. We denote the location of CRMs-TFs complex on DNA as *X*. CRMs spans a length *X* = (0, *X*_*0*_) and the promoter is located at *X* = *L*. **B**. Looping mediated propulsion of combinatorial TFs complex with the overall radius of gyration of *r*_*P*_ bp along DNA. In this example, there are five different TFs (denoted as TF-1, 2, 3, 4, 5 respectively) involved in the combinatorial regulation. Here DBD1 type DBD of TF-1 specifically binds at S1 (CRMs) and DBD2 of TF-4 specifically binds at the promoter. TFs 2, 3, and 5 involve in the stabilization of the CRMs-TFs complex via protein-protein interactions. TF-1 which arrives first at S1 will further recruit other four TFs in a cooperative manner. This complete CRMs-TFs complex is stabilized by the site-specific hydrogen bonding interactions between DBD1 of TF-1 and the CRMs located at S1 of DNA, protein-protein interactions among all the five TFs and the cumulative nonspecific electrostatic interactions between DBD2s of other TFs (2, 3, 4 and 5) with DNA. Binding of these TFs near CRMs (that spans for a length of *X*_0_ from S1 (*X* = 0) to *X* = *X*_0_) bends the DNA segment into a loop around them such that *X* _0_ = 2π *r*_*P*_. The elastic stress on DNA can be incrementally released via bulging of DNA around CRMs-TFs complex. **B**. Since the specific binding of DBD1 of TF-1 at S1 as well as the cumulative nonspecific interactions of all the TFs are strong enough to resist the dissociation, the elastic stress of DNA in the CRMs-TFs complex can be released only via thermally induced sliding of DBD2 of TFs towards the promoter well within the Onsager radius associated with the DBD2s-DNA interface. Upon reaching the promoter, DBD2 of TF-4 interacts with the promoter to form the final synaptosome complex. **D**. Synaptosome complex where the combinatorial TFs are bound with both CRMs and the promoter via DNA-loop.

### Energetics of the site-specific binding of TFs and bending of DNA

Let us assume that the radius of gyration of the fully assembled combinatorial TFs complex is *r*_*P*_ bp (Fig. 1B) and CRMs are located at S1 of DNA. We further assume that there are two different type of DBDs of TFs viz. DBD1 and DBD2. Here DBD1 site-specifically binds CRMs via hydrogen bonding and DBD2s bind near CRMs via nonspecific electrostatic interactions. We denote the distance between CRMs and the promoter by the variable *X* (measured in bp). Upon binding the cognate stretch of DNA with size of *X*_*0*_ bp, the TFs complex bends the DNA segment into a circle around its overall spherical solvent shell surface such that the radius of curvature of the bent DNA segment is same as that of the radius of gyration of the combinatorial TFs complex. In this situation *X* _0_ ≃ 2π *r*_*P*_ as shown in Fig. 1B. In general, one can write as *X*_0_ ≃ δ*r*_*P*_ where 0 < δ ≤ 2π. We set *X* = 0 at S1 and *X* = *X*_0_ at the end of the DBD2s of TFs on DNA. Here S1 is the specific site for DBD1 and P is the specific site for DBD2s by definition. Here segment of DNA under consideration spans over the range (0, *L*) as in Fig. 1A and *X* is the current loop-length. The total energy required to bend a linear piece of DNA will be the sum *E*_bend_ = *E*_elastic_ + *E*_entropy_ and in general *E*_bend_ ≥ 0. For the radius of curvature *r*_*P*_, one finds that 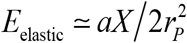 (*kBT* units) where *α* is the persistence length of DNA (37, 50). Clearly, *E*_elastic_ that is required to bend a DNA segment of length *X* into a circle will be *E*_elastic_ ≃ 2π^2^*a/X*. Here one can write *E*_elastic_ ≃ δ^2^*a*/2*X* where 0 < δ ≤ 2π. Clearly, *E*_elastic_ will be at maximum when δ = 2π. This energy has to be derived either solely from the binding of TFs on DNA or via an external energy input in the form of ATP hydrolysis (51). Noting that *E*_entropy_ ≃ (3/2) ln(*πX*/6*b*) where *b* = 1 bp is the distance between two consecutive nucleotide monomers of DNA, one finally arrives at the following expression for the overall bending energy (**Eq. A1** of **Appendix A**).

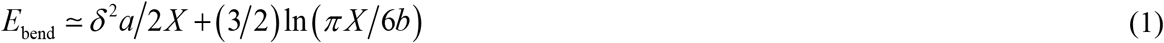

Clearly, *E*_bend_ attains a minimum value as 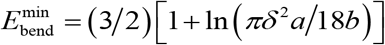 *X*_*C*_ = δ^2^*a*/3. In the later sections, we will show that this non-monotonic behavior of the bending energy profile will restrict the possible distances between CRMs and their corresponding promoters.

### Looping mediated communication between CRMs-TFs and promoters

Let us assume that the first arrived TF at CRMs recruits other combinatorial TFs via protein-protein interactions. When all the TFs bind their CRMs in the first step of transcription activation and bend the DNA over their spherical solvent shell surface, then the overall *binding energy* (*E*_bind_) can be written as *E*_bind_ = *E*_bend_ +; *E*_elastic_ + *E*_entropy_. In general, one should note that *E*_bind_ ≤ 0 and *E*_elastic_ ≥ 0. Apparently, a stable complex of CRMs-TFs can be formed only when *E*_bind_ < 0. The bonding energy released at the CRMs-TFs interface is utilized to compensate both the elastic energy required to bend the DNA (*E*_elastic_) and the chain entropy loss (*E*_entropy_) at the specific binding site. One should note that *E* ≃ |*E*_bond_| + *E*_elastic_ is the overall potential energy barrier which acts against any kind of distortion or dissociation of the CRMs-TFs complex. Here *E*_bond_ comprises of energies associated with the formation of site-specific hydrogen bonding network (DBD1-S1 interactions), protein-protein interactions among the combinatorial TFs and the overall cumulative non-specific interactions between the DBD2s of TFs and DNA i.e. *E*_b0nd_ = *E*_hydrogen_ + *E*_protein-protein_ + *E*_electrostatic_. Among these three, only the electrostatic interactions can be easily perturbed by the thermally induced fluctuations. Conversely, *E* ≃ *E*_elastic_ + *E*_entropy_ is the potential energy barrier that resists the formation of loops out of a linear piece of DNA. With this background, the elastic stress involved in the bent DNA of CRMs-TFs complex can be released via the following four different modes (types I, II, III and IV) of dynamics depending on the number of TFs (*n*) involved in the combinatorial regulation. For the comparison purpose, we denote the number of TFs involved in these types of dynamics as *n* = (*n*_*I*_, *n*_*II*_, *n*_*III*_, and *n*_*IV*_).

I. Thermally induced physical dissociation of the entire CRMs-TFs complex along with increase in the overall chain entropy. This mechanism will be frequent especially when *n*_*I*_ = 1. However, when the number of TFs involved in the nonspecific electrostatic interactions is high, then their cumulative electrostatic interactions will be strong enough to resist the thermally induced dissociation of DBD2s of the CRMs-TFs complex from DNA. Clearly, when *n* is sufficiently large then the thermally induced spontaneous dissociation of the CRMs-TFs complex is not possible in the physiologically relevant timescales.
II. Physical dissociation of only DBD2s of TFs complex from DNA and their re-association somewhere else via looping over 3D space (which is resisted by the loop-length dependent potential energy barrier *E* ≃ *E*_elastic_ + *E*_entropy_) while the specific interactions at S1-DBD1 of main TF is still intact. This is the *tethered binding-unbinding model* of Shvets and Kolomeisky developed in Ref. (52). In this model, there is no restriction on the initial value of *X* i.e. the DBD2s of TFs can land anywhere within (0, *L*). Additionally, the sliding of DBD2s on DNA is not allowed in this model. This model will work only when the nonspecific electrostatic interactions associated with the DBD2s of TFs are weak enough to dissociate frequently. This in turn necessitates a small number of combinatorial TFs.
III. When the number of TFs involved in the nonspecific electrostatic interactions is high enough, then the thermally induced physical dissociation of DBD2s of TFs will be less probable. However, thermally induced sliding of DBD2s is still possible well within the *Onsager radius* associated with the nonspecifically bound regions of DBD2s-DNA interface. Since there is a reflecting boundary condition imposed by the strong DBD1-S1 interactions at CRMs, there is a possibility for the directional-dependent propulsion of the CRMs-TFs complex on DNA via sliding of DBD2s of CRMs-TFs complex towards the promoter. This is achieved through gradual increase in the value of *X* from *X*_*0*_ towards *L* while the specific interactions at S1-DBD1 of main TF is still intact. This is the *directional-dependent propulsion model*.
IV. Tethered sliding of DBD2s of TFs with dynamic loop structure of DNA and intact DBD1-S1 interactions at CRMs. This is similar to type II where sliding of DBD2s of TFs on DNA is allowed before unbinding. This further necessitates a small number of combinatorial TFs (lesser than type III but more than II). Depending on the number of combinatorial TFs involved in the regulation one can consider two different possibilities here viz. a *pure tethered sliding* and *tethered binding-sliding-unbinding*. In pure tethered sliding, the DBD2s of the CRMs-TFs complex forms nonspecific contacts at an arbitrary location on DNA via a dynamic loop structure and then searches for the promoter region via sliding dynamics without dissociation. Here the cumulative electrostatic interactions between the DBD2s and DNA are strong enough to retain the dynamic loop structure until DBD2s reach the promoter. In the tethered binding-sliding-unbinding mode, the DBD2s of CRMs-TFs performs several cycles of binding-sliding-unbinding before finding the promoter region.

One can arrange the number combinatorial TFs involved in II, III and IV types as *n*_*II*_ < *n*_*IV*_ < *n*_*III*_. In the propulsion mechanism, mainly a gradual release of the elastic stress increases the radius of curvature of the bent DNA. This in turn causes bulging of the DNA-loop around the TFs complex as described in Fig. 1C. The chain entropy does not increase much here since the intervening DNA is still under loop conformation. This is similar to the sliding of nucleosomes via bulge induced reptation dynamics of DNA (14–16). In general, dissociation of CRMs-TFs complex will be an endothermic process since |*E*_bond_| > |*E*_elastic_|. *Clearly*, *when the number of combinatorial TFs is large enough*, *spontaneous physical dissociation of the CRMs-TFs complex will not be the most probable route for the release of elastic stress of the bent DNA polymer*. With this background, the DBD2s of TFs complex need to distally interact with the promoter in the second step and activate the transcription via looping of the intervening DNA segment that connects CRMs and the promoter. In this context, only II, III and IV types of dynamics can be considered here and we rule out the possibility of the type-I dynamics. The repeated binding-unbinding model that is characterized by the type II dynamics of CRMs-TFs complex has been studied in detail by Shvets and Kolomeisky (52). Actually, type II is a special case of type IV dynamics. In the following sections we will consider the possibility of types III and IV in the looping mediated transcription activation in detail i.e. directional-dependent propulsion and tethered sliding models. All the symbols used in this paper are listed in Table S1 of the Supporting Material.

### Directional-dependent propulsion model

The main idea of this model is as follows. Let us assume that the binding energy profile of the combinatorial TFs is such that the specific binding near DBD1-S1 as well as the cumulative nonspecific electrostatic interactions between DBD2s and DNA are strong enough to resist the abrupt dissociation of TFs from DNA for prolonged timescales. In this situation, since the DBD1-S1 is a strong specific binding (reflecting boundary), the elastic stress of the bent DNA of the CRMs-TFs complex can be released only via the thermally driven sliding of DBD2s well within the Onsager radius associated with the DBD2s-DNA interface. When the DBD2s of CRMs-TFs complex slide along DNA, then the elastic stress on DNA will be released in a gradual manner via bulging of the DNA-loop around the CRMs-TFs complex. This in turn propels the CRMs-TFs complex towards the promoter located at *L* as shown in Figs. 1B-D. A schematic representation of the propulsion model is given in Fig. S1 of the Supporting Materials. Here the water molecules present at the interface of the positively charged DBD2s of TFs and the negatively charged backbone of DNA provide a fluidic type environment for the smooth sliding dynamics of CRMs-TFs complex (8, 21). Upon finding the promoter, CRMs-TFs forms tight synaptosome complex that is required for the transcription activation. Clearly, the following two conditions are essential for the feasibility of propulsion model.

a. The binding energies associated with the specific and nonspecific interactions should be strong enough to bend the CRMs-DNA segment around the TFs complex.
b. The overall cumulative nonspecific binding energies associated with the DBD2s-DNA interactions should be strong enough to sustain the sliding of DBD2s with intact dynamic loop structure of DNA until finding the promoter.

Using detailed calculations, we will show in the later sections that both these conditions can be fulfilled by raising the number of TFs involved in the combinatorial regulation. Although there is no straightforward experimental evidence for this model, one can construe this idea indirectly from various other experimental studies. Particularly, Rippe et.al (32) have studied NtrC (Nitrogen regulatory protein C) system using the scanning force microscopy. In this study, they had taken snapshots of various intermediary states along the process of transcription activation from the closed to the open promoter complex. In their model system, binding of NtrC at its specific site (CRM) activates the downstream closed complex of *glnA* promoter-RNAP-σ^54^ via looping out of the intervening DNA segment. They have shown that the transition from the inactive-closed form to an active-open promoter complex involved a gradual increase in the bending angle of the intervening DNA. This in turn is positively correlated with an increase in the radius of curvature of the intervening DNA segment which is represented as bulging of the DNA-loop in our propulsion model. Therefore, our assumption that the propulsion of TFs via increase in the radius of curvature of the bent DNA is a logical one. Based on these, the dynamical position *X* of TF on DNA obeys the following Langevin type stochastic differential equation (53–55).

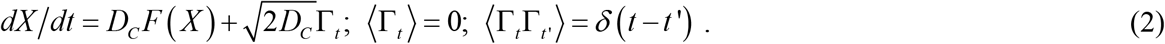

In Eq. 2, *F* (*X*)= −*dE*/*dX* = 2π^2^*a*/*X*^2^ (bp^−1^) is the force acting on DNA of the CRMs-TFs that is generated by the potential *E* ~ *E*_elastic_ + |*E*_bond_| upon bulging of the loop structure, Г, is the Δ-correlated Gaussian white noise and *D*_*C*_ (bp^2^/s) is the 1D diffusion coefficient of the sliding of TFs. The energy involved in the bonding interactions will be a constant one so that it will not contribute to the force term. We ignore the energy dissipation via chain entropy of the bulging loop structure mainly because binding of TFs at CRMs attenuates the conformational fluctuations at the CRMs-TFs interface (8, 20, 21). The Fokker-Planck equation describing the probability of observing a given *X* at time *t* with the condition that *X* = *X*_0_ at *t* = *t*_0_ can be written as follows (53, 54).

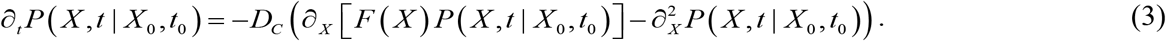

The form of *F*(*X*) suggests that it can propel the DBD2s of CRMs-TFs complex only for short distances since lim_*X* →∞_ *F* (*X*) = 0 although such limit will be meaningless for *X* > 2*π*^2^*a* where *E*_elastic_ will be close to the background thermal energy. Further, one also should note the fact that the force *F*(*X*) changes sign below some critical value of *X* (56). Initial condition for Eq. 3 will be *P* (*X*, *t*_0_ | *X*_0_, *t*_0_) = δ(*X* −*X*_0_) where *X* _0_ = 2π r_*p*_ and the boundary conditions are given as follows.

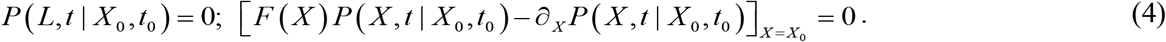

Here *X*_0_ acts as a reflecting boundary for a given size of TFs complex and *L* is the absorbing boundary where promoter is located. The strong site-specific interactions at the DBD1-S1 of TFs complex act as a reflecting boundary condition. The mean first passage time (MFPT) *T*_*B*_ (*X*) associated with the DBD2s of CRMs-TFs to reach the promoter location *L* starting from an arbitrary *X* ∈ (*X*_0_, *L*) obeys the following backward type Fokker-Planck equation along with the appropriate boundary conditions (7, 8).

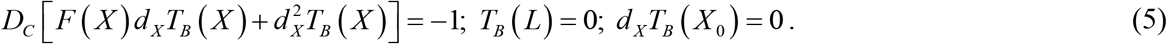

The integral solution of Eqs. 5 can be expressed as follows.

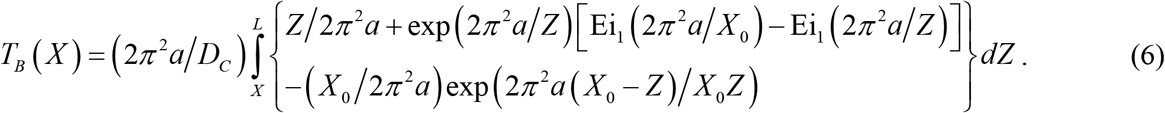

Here 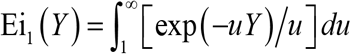(57) and interestingly lim_*L*→∞_ *T*_*B*_(*X*). Here *T*_*N*_(*X*) is the mean first passage time required by the DBD2s of TFs to reach *L* via pure 1D sliding in the absence of dynamic loop structures of DNA which is a solution of the following differential equation (7, 8, 21).

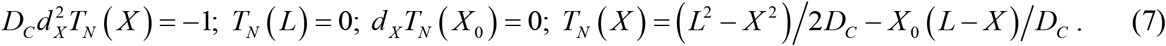

To obtain the target finding time, one needs to set *X* = *X*_0_ in Eqs. 6 and 7. One can define the number of times the target finding rate of TFs can be accelerated by the looping mediated propulsion of TFs over 1D sliding as 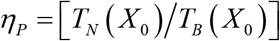(here the subscript ‘*P*’ denotes the propulsion model) which is clearly independent of *D*_*C*_ of TFs complex and solely depends on (*L*, *a*, and *X*_*0*_). Explicitly one can write it as,

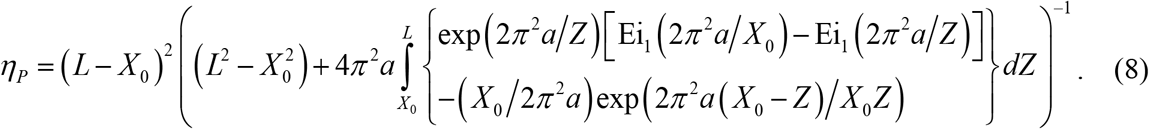

Detailed numerical analysis (Section 2 of the **Supporting Material**) suggests that there exists a maximum of η_*P*_ at which ∂η_*P*_/∂*L* = 0 with *L* = *L*_opt_ and clearly, we have lim_*L*→∞_ η_*P*_ = 1 (Figs. 2A and B). This is logical since when *L* > *L*_opt_ then η_*P*_ → 1 and when *L* < *L*_opt_ then the stored energy is not completely utilized to propel the DBD2s of CRMs-TFs. Further, 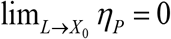 since its numerator part goes to zero much faster than the denominator part (Figs. S2 and S3). The total time required by the combinatorial TFs to form a synaptosome complex via propulsion mechanism will be τ_*p*_ = τ_*s*_ + *T*_*B*_(*X*).

**FIGURE 2.**
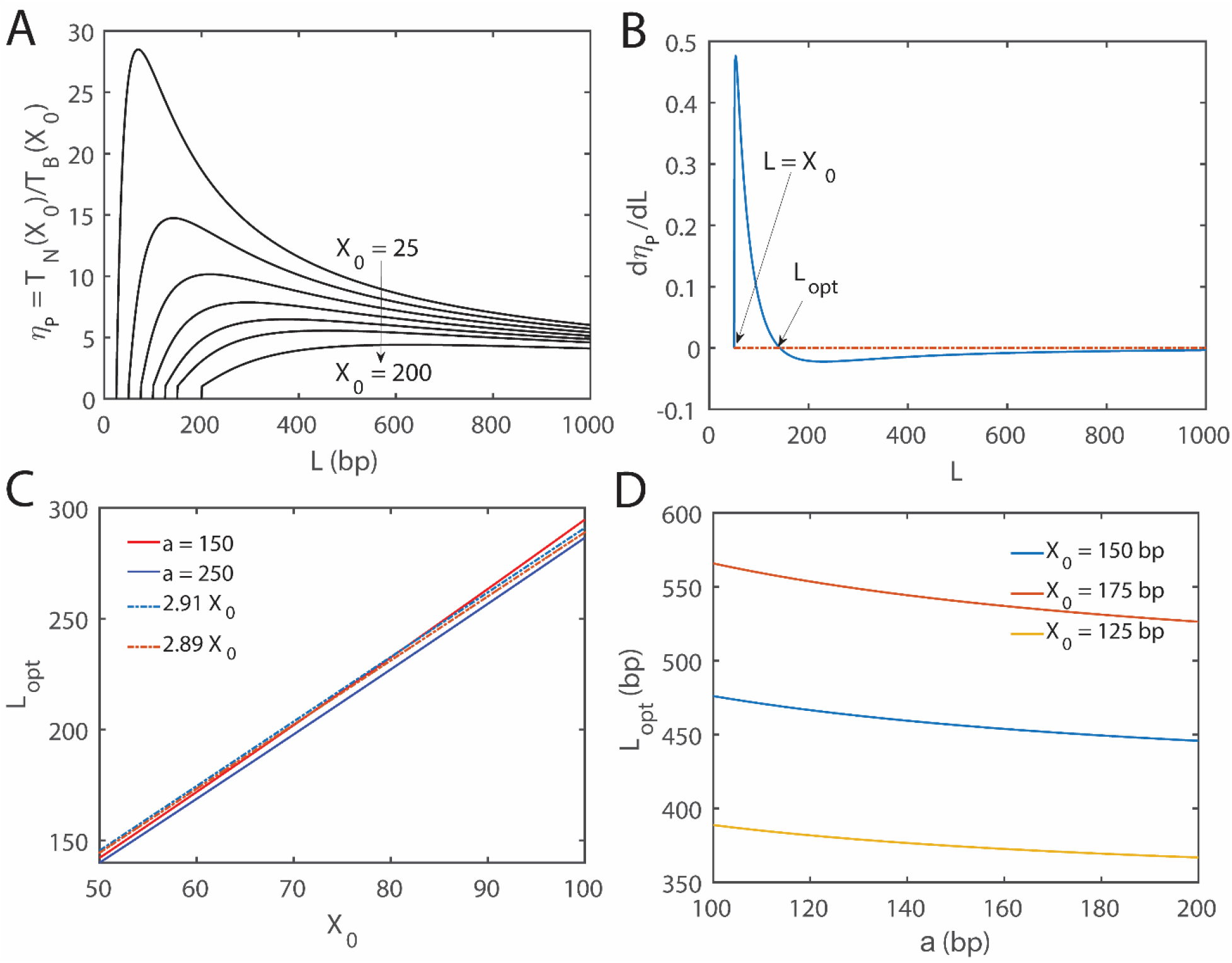
Numerical analysis of the propulsion model. **A.** Relative efficiency of looping mediated stochastic propulsion of TFs versus normal 1D sliding along DNA. *T*_*N*_(*X*_0_) is the mean first passage time that is required by TFs to reach the promoter that is located at *L*, starting from *X*_0_ via 1D sliding. *T*_B_(*X*_0_) is the mean first passage time required by TFs to reach *L* starting from *X*_0_ via looping mediated stochastic propulsion mechanism. *X*_0_ was iterated as (25, 50, 75, 100, 125, 150, 200) along the arrow while iterating *L* from *X*_0_ to 1000. The efficiency of looping mediated sliding is strongly dependent on the persistence length of DNA (*a*), *L* and *X*_0_ and it is a maximum at *L*_opt_ ~ 3*X*_0_. **B**. Plot of *dη*_*P*_/*dL* with respect to *L*. Here the settings are *a* ~ 150 bp and *X*_0_ ~ 50 bp and *L* was iterated from 50 to 1000 bp. Upon solving *dη*_*P*_/*dL* = 0 for *L* numerically one finds that *L*_opt_ ~ 142.2 bp. **C**. Variation of *L*_opt_ with respect to *X*_0_. Clearly *L*_opt_ ~ 3*X*_0_, is slightly dependent on the persistent length *a*. Here we have iterated *X*_0_ from 50 to 100 bp and *a* = (150, 250) bp. The solution for *L* was searched within the interval (50, 1000) bp. **D**. Variation of *L*_opt_ with respect to changes in *a*. Here we have iterated *a* from 100 to 200 bp and *X*_0_ = (125, 150, 200) bp. The solution for *L* was searched within the interval (50, 1000) bp. The error in the approximation *L*_opt_ ~ 3*X*_0_ seems to be < 10% over wide range of *a* values.

### Predictions of the propulsion model

The persistence length of DNA under *in vitro* conditions is *a* ~ 150 bp and the radius of gyration for most of the eukaryotic TFs will be around ~ 5 bp. When the number of TFs involved in the combinatorial regulation is 3-5, then the maximum *r*_*P*_ ~ 15-25 bp. Therefore, one can set the initial *X* = 2π *r*_*P*_ ~100-150 bp where we have set *δ* = 2*π* here (58, 59). Simulations (Fig. 2A) of the expression for η_*P*_(Eq. 7) at different values of *X_0_*and, *L* from *X_0_* to 10^5^ suggested that *L*_opt_ ~ 3*X*_0_ (Figs. 2C and 2D). When *a* ~150 bp and *X*_0_ ~ 100-150 bp, then *L*_opt_ ~ 300-450 bp. Remarkably, this is the most probable range of the distances between the CRMs and promoters of various genes observed across several genomes (60). The efficiency of the propulsion will be maximum at *L*_opt_. Although *L*_opt_ is not much affected by *a*, the maximum of *η*_*P*_ is positively correlated with *a*. This is logical since the stored elastic energy is directly proportional to the persistence length of the polymer. Remarkably, at the optimum *L*_opt_ the speed of interactions between CRM-TFs complex with the promoters will be ~10-25 times faster than the normal 1D sliding.

### Tethered sliding models

When the number of TFs involved in the combinatorial regulation is less such that *n*_*IV*_ < *n*_*III*_, then the cumulative nonspecific electrostatic interactions will not be enough to bend the DNA around the TFs complex as in the case of propulsion model (Fig. 3). Therefore, stable initial nonspecific contacts between DBD2s and DNA will be formed slightly away from CRMs (S1) i.e. the radius of curvature of the initial looped-out DNA structure will be higher than the radius of gyration of the TFs complex *r*_*P*_. Here the tethered random walker (DBD2s of CRMs-TFs complex, which is actually tied with the DNA thread at DBD1-S1) wanders over 3D space and randomly forms nonspecific contacts with other segments of the same DNA polymer analogous to the ring-closure events of intersegmental transfer dynamics. Before dissociation, DBD2s of CRMs-TFs may scan the DNA of random length for the presence of promoter P. In pure tethered sliding, the nonspecific electrostatic interactions between DBD2s and DNA are strong enough to keep the dynamic loop structure intact until reaching the promoter via sliding dynamics. In case of tethered binding-sliding-unbinding, the tied DBD2s perform multiple cycles of binding-sliding-unbinding before reaching the promoter.

**FIGURE 3.**
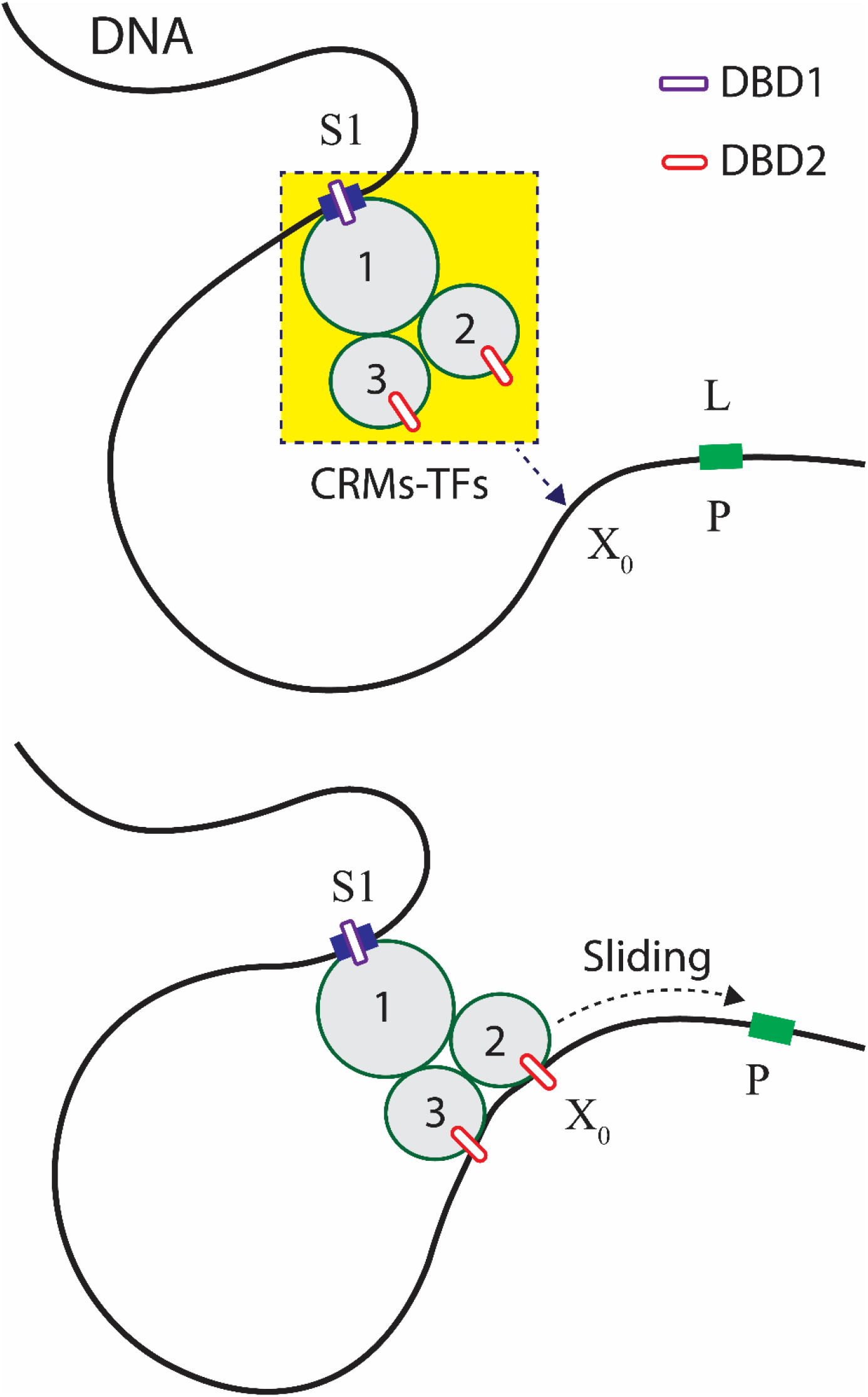
Tethered sliding model. In this example, there are three combinatorial TFs involved in the regulation viz. TF-1, 2 and 3. Binding of TF-1 at the CRM located at S1 subsequently recruits TF-2 and 3 near the location of S1 via protein-protein interactions. The DBD2s of TF-2 and 3 are specific to the promoter located at P whereas the DBD1 of TF-1 is specific to the CRMs located at S1. With intact hydrogen bonding interactions at S1, the CRMs-TFs complex wander over 3D space and forms nonspecific contacts with DNA randomly at *X*_0_. Depending on the strength of the cumulative nonspecific interactions of DBD2s, one can consider pure tethered sliding or tethered binding-sliding-unbinding. In pure tethered sliding, upon forming the first nonspecific contact with DNA at *X*_0_ via dynamic loop structure, DBD2s of TFs search for the promoter via sliding without dissociation. In case of tethered binding-sliding-unbinding, nonspecifically bound DBD2s of TFs perform several cycles of nonspecific binding-sliding-dissociation before finding the promoter. When the DBD2s of TFs find the promoter that is located at P, then it forms the final synaptosome type complex.

We first consider the pure tethered sliding model. When the length of DNA that connects DBD1-S1 and the landing position of DBD2s is *X* for an arbitrary nonspecific contact of DBD2s, then the potential energy barrier acting on such random sliding will be *E* ≃(2π^2^*a*/*X*) + (3/2) ln (π*X*/6*b*) where *b* = 1 bp is the distance between two consecutive nucleotides of DNA. This potential energy barrier attains a minimum value as *E*_min_ = [3/2](1+ ln (2π^3^*a*/9*b*)) at *X*_C_ = 4π^2^*a*/3. Forward and reverse movement of such *tethered random walker* drives *X* to *X* + 1 or *X* − 1. Contrasting from the propulsion model, here we have not ignored the entropy component of the potential function *E* since the initial interconnecting DNA segment is larger than the radius of gyration of the TFs complex and also in the free loop form. As a result, one cannot ignore the entropic barrier associated with the loop formation. Moreover, sliding of CRMs-TFs with dynamic loop structure will always be impeded by the chain entropy. The force generated by such potential will be *F* (*X*)= *X* 2π^2^*a*/*X*^2^ − 3/2*X*. Upon inserting this force term into Eq. 5 one finally obtains the following integral solution.

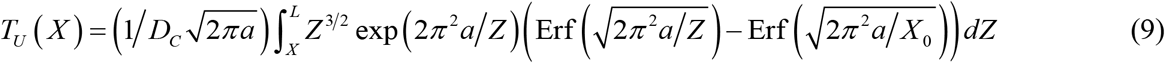

Here 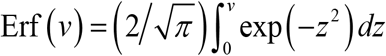 is the error function integral (57), and *T*_*U*_(*X*) is the MFPT required by a pure tethered random walker to find its specific site located at *L* starting from *X* (this is the initial loop length) anywhere within (*X*_0_, *L*) where *X*_0_ is a reflecting boundary and *L* is an absorbing boundary. Since the potential function has a minimum at *X*_*C*_, one can consider the following two different limiting regimes.

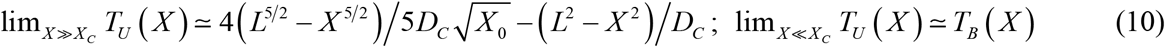

One can define the number of times the target finding rate of TFs can be accelerated by the tethered sliding of TF as η_S_ = *T*_*N*_ (*X*)/*T*_*U*_(*X*) (here the subscript ‘*S*’ denotes the tethered sliding model) which is clearly independent of *DC* of TF and solely depends on (*L*, *a*, and *X*_0_). Contrasting from the propulsion model, one finds that lim_*L*→∞_η_*S*_ = 0. In these calculations we have not included the looping mediated nonspecific association time required by the DBD2s of the CRMs-TFs complex. This in fact further increases the overall MFPT of the tethered sliding model. The rate associated with the formation of the initial (nonspecific contact) loop with contour length *X* can be written as *k*_*NL*_ ≃ *k*_*t*_ exp (−*E*) where *k*_*t*_(*s*^−1^) is the maximum achievable rate under zero potential. Clearly, *k*_*NL*_ will be a maximum at *X*_*C*_ which is the most probable initial landing position of the tethered DBD2s of TFs via DNA-looping. In this model, the total time required by the CRMs-TFs to form the final synaptosome complex will be τ_*TS*_ =τ_*S*_ + 1/*k*_*NL*_ + *T*_*U*_(*X*) which attains a local minimum near *X* = *X*_*C*_. One can also define η_*NL*_ = *k*_*NL*_/*k*_*t*_ which attains the maximum η_*NL*_ ~ 6.7 *X*_*C*_. Since there are several unbinding events in the tethered binding-sliding-unbinding model, the overall time required by the DBDs to form the synaptosome complex will be much higher than τ_*TS*_.

### Predictions of the tethered sliding models

Tethered sliding model predicts the most probable distance of the CRMs of TFs from the transcription start sites as *X*_*C*_. At this distance, the rate of looping mediated synaptosome complex formation of TFs will be at maximum. Upon setting *X* = *X*_*C*_ in *η*_*S*_ and numerically iterating *L* from 3000 to 10000 bp with *a* ~ 150 bp, one can observe the following results. When the left reflecting boundary was at *X*_*0*_, then one finds the critical distance *L*_*C*_ such that *η*_*S*_ > 1 when *L* < *L*_*C*_ and approximately *η*_*S*_ < 1 when *L* > *L*_*C*_. Particularly, when *X*_0_ < 100 bp then one can define the critical distance of transcription start sites from CRMs in the tethered sliding model as *L*_*C*_ ~ 3*X*_*C*_. This critical distance decreases with increase in *X*_*0*_. These numerical results are demonstrated in Fig. S4 of the Supporting Materials.

## COMPUTATIONAL ANALYSIS

Tethered sliding model predicted the most probable distance of CRMs from the transcription start site (promoter, location S1 on DNA) as *X*_*C*_ = 4π^2^*a*/3 ~ 2000 bp for *a* ~ 150 bp. To check the validity of this prediction, we analyzed the upstream 5000 bp sequences of various genes of human and mouse genome. We used the position weight matrices of various transcription factors of human and mouse available with the JASPAR database and scanned upstream sequences of all the genes of the respective genomes.

### Datasets and analysis

The upstream 5000 bps sequences of various genes of human and mouse genomes were obtained from UCSC genome database (February 2009 assembly, hg19 version for human genome and December 2011 assembly, mm10 version of mouse genome) and position weight matrices (PWMs) (61, 62) of various TFs of mouse and human were obtained from the publicly available JASPAR database (63, 64). There were 21929 sequences from mouse genome and 28824 sequences from the human genome. Using the PWMs of various available TFs we generated the score table for various upstream sequences based on the following equation (61).

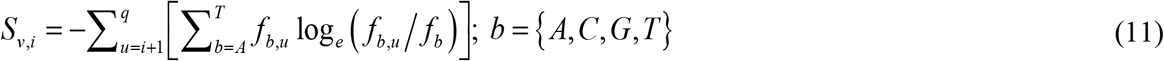

In this equation *S*_*v,i*_ is the score value of PWM at *i*^th^ position upstream of the transcription start site (TSS) on *v*^th^ sequence, *q* is the length of binding stretch of the corresponding TF, *f*_*b*_ is the background probability of observing base *b* in the corresponding genome, and *f*_*b,w*_ is the probability of observing base *b* at position *w* of the specific binding sites of TFs. Here *f*_*b*_ was calculated from the random sequences of the given genome available with the UCSC database. We constructed the distribution of the distances of S1 from the transcription start site. There is a strong positive correlation between the score value and the binding energy of TFs (61). Computing the distribution of the distances of S1 from the TSS will confirm the validity of the tethered sliding model.

In parallel, we also generated score table for random sequences using the same PWM from which we obtained the score distribution and the cutoff score value for the given weight matrix corresponding to a given *p*-value. In our calculations, we have set the *p*-value < 10^−6^ for defining the putative specific binding sites of TFs. We used the random sequences associated with each genome that is available at UCSC database to compute the probability of occurrence of putative binding sites by chance. We considered random sequences of size 5 × 10^6^ bps and fragmented it into 10^3^ number of sequences with length of 5000 bps. Then we scanned each random sequence with the same PWM and obtained the number of putative CRMs (false positives). The probability of observing a CRM site by chance will be calculated as *p*_*NF*_ = number of false positives / 1000.

## RESULTS AND DISCUSSION

Various possible modes of transcription activation via combinatorial binding of several TFs are summarized in Fig. S5 of the Supporting Materials. The main limitation of the propulsion model is the requirement of huge energy input involved in the initial bending of DNA around the TFs complex of interest. This needs to be derived either in the form of ATP hydrolysis or in the form of binding energy derived from the combinatorial TFs. For example, bending of a linear DNA with size of 100-150 bp into loop requires the hydrolysis of at least 2-3 ATPs (using *E*_bend_ = *E*_elastic_ + *E*_entropy_, here hydrolysis of 1 ATP molecule releases ~12 *k*_*B*_*T*). Investment of such energy input is required by CRMs-TFs system to actively slide in a directional dependent manner towards the promoter. Instead, tethered sliding of TFs does not require such huge energy input since there is no restriction on the initial loop length. As a result, directional dependent movement of TFs is not possible in the tethered sliding model. However, the probability density function associated with the initial loop length will be dictated by the bending energy profile. Actually, *E*_bend_ will be a minimum at *X*_*C*_ ≃ 4π^2^*a*/3 where the average search time required to form the final synaptosome complex will be at minimum (52). When *X* < *X*_*C*_ then *E*_bend_ ∝ *X*^−1^. When *X* > *X*_*c*_ then one finds that *E*_bend_ ∝ ln (*X*). When *a* ~ 150 bp and *X*_*C*_ ~ 2000 bp then the minimum of *E*_bend_ ~ 13 *k*_*B*_*T* which requires the hydrolysis of at least 1 ATP. These results are depicted in Fig. 4.

**FIGURE 4.**
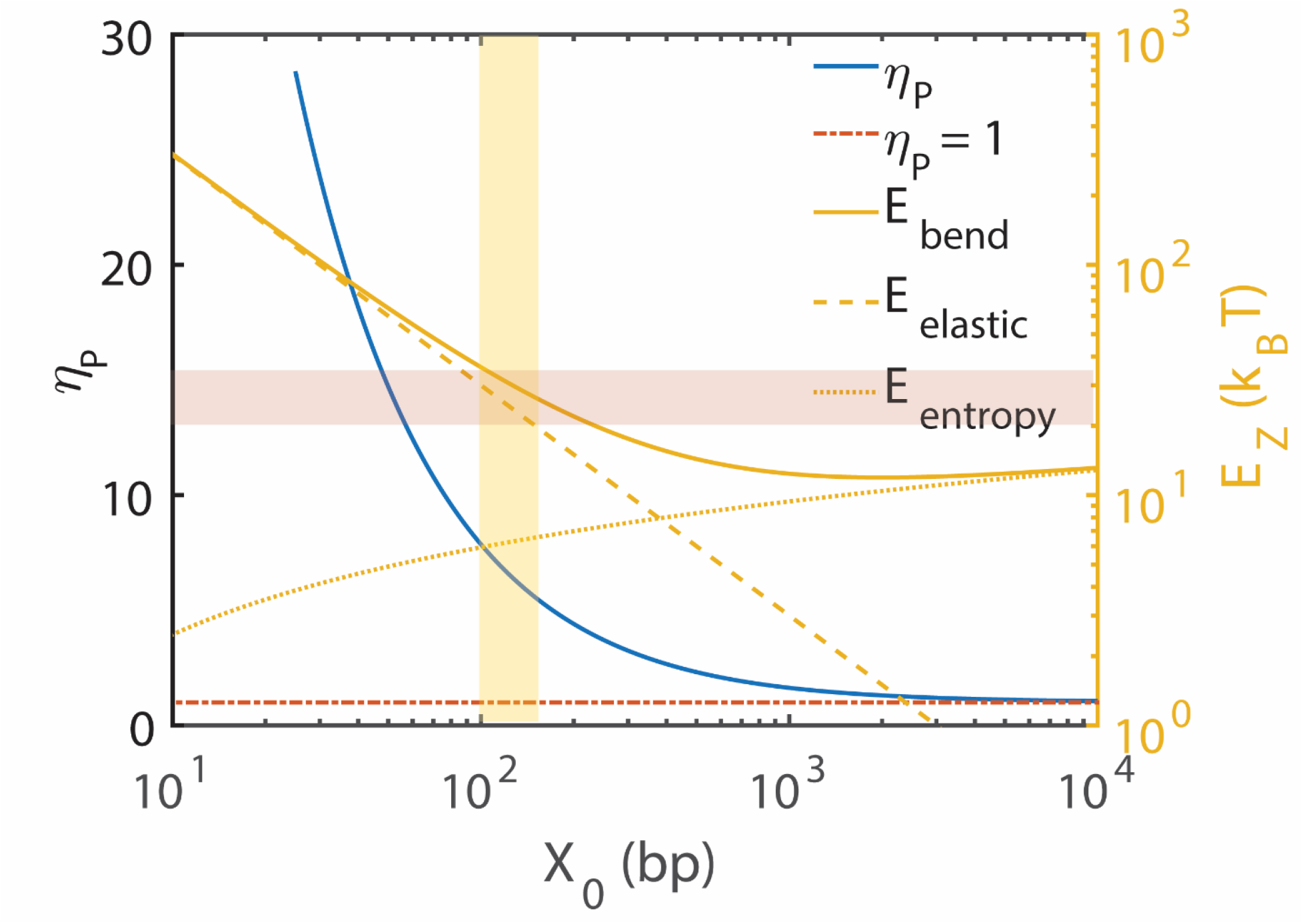
Energy components of looping mediated transcription activation. Variation of the propulsion efficiency *η*_*P*_ and bending energy with respect to changes in the initial loop length *X*_0_. Here the settings are *a* ~ 150 bp, *L* = 3*X*_0_ ~ *L*_opt_. We computed η_*p*_ = *T*_*N*_ (*X*_0_)/*T*_*B*_(*X*_0_) with *L* = 3*X*_0_ so that *η*_*P*_ will be close to its maximum. Here the subscript *Z* = (entropy, bend, elastic, enthalpy). *E*_bend_ = *E*_elastic_ + *E*_entropy_ where *E*_elastic_ ≃ 3000/*X*_0_ and *E*_entropy_ ≃ 3 ln (π*X*_0_/6*b*/2 which is ~12 *k*_*B*_*T* at *X*_0_ ~ 2000 bp. *E*_elastic_ ≤ 1 *k*_*B*_*T* when *X*_0_ ≥ 3000 bp. Clearly, the bending energy of the linear DNA is always ≥ 12 *k*_*B*_*T* irrespective of the length. Shaded regions are the most probable *X*_0_ values observed in the natural systems (~100-150 bp). The corresponding optimum distance between the transcription factor binding sites and promoters *L*_opt_ ~ 3*X*_0_ ~ 300-450 bp.

The analysis results on the upstream sequences of various genes of human and mouse are shown in Figs. 5. The most probable location of S1 seems to be around ~2500 bp away from the promoter in both mouse and human genome. The distributions of the distances of CRMs from the respective TSS are shown in Figs. 5A and B. These results are in line with the tethered sliding model which predicted the critical distance of CRMs from the promoter to be around *X*_*C*_ ~ 2000 bp. Including various models presented here, one can consider the following four possible modes associated with the formation of synaptosome complex.

**FIGURE 5.**
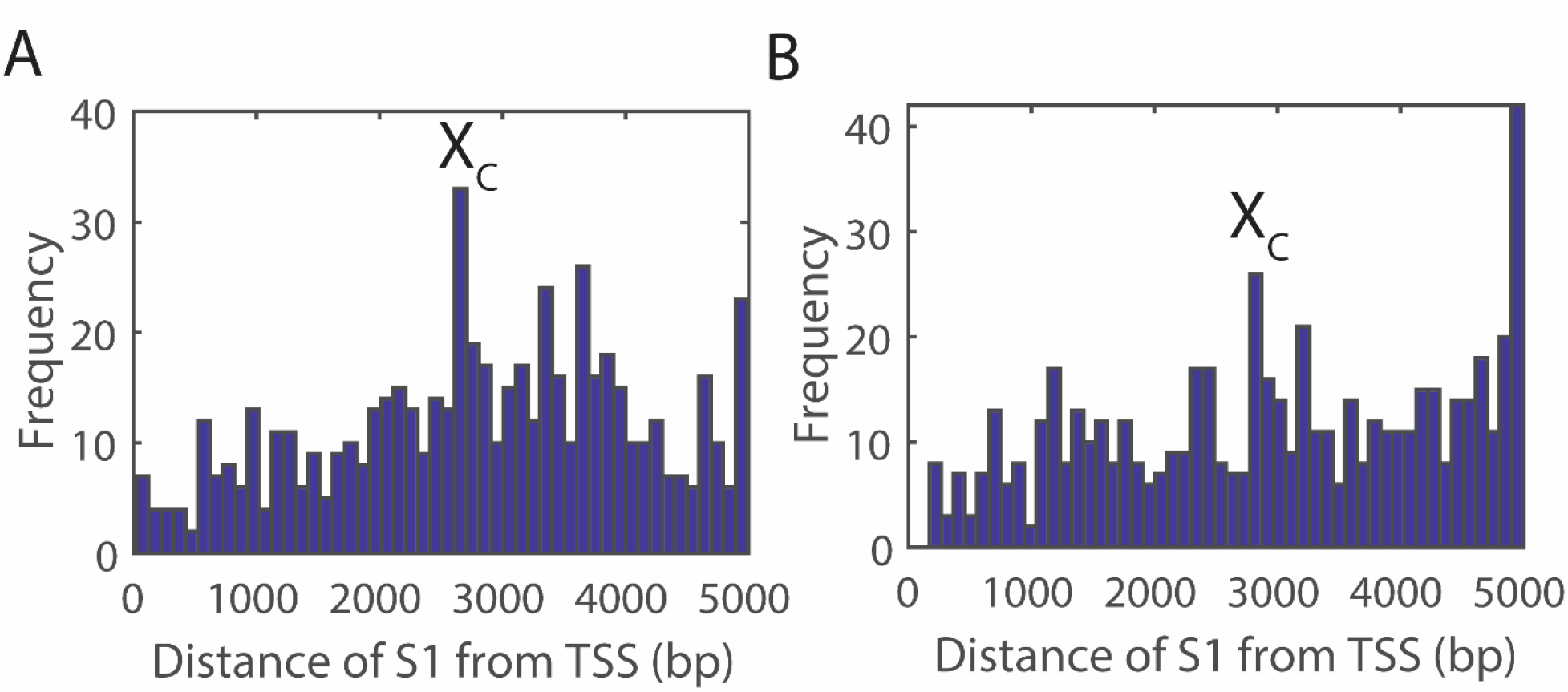
Computational data analysis results. We considered the position weight matrices of human and mouse available with the JASPAR database and scanned the upstream 5000 bp sequences of various human and mouse genes. The tethered sliding model predicted the most probable distance of the CRMs of TFs from the transcription start sites as *X*_*C*_ ~ 2000 bp for a persistence length of DNA as *a* ~ 150 bp. The distribution of the upstream location of CRMs of various TFs shows a maximum around ~2500 bp. Putative binding sites were defined with a *p*-value < 10^−6^. These results are in line with the tethered sliding model. **A**. Mouse. **B**. Human.

a. *Propulsion mechanism*. This requires huge free energy input in the initial loop formation with a possibility of directional dependent movement of TFs towards the promoter. Eventually this mechanism warrants several combinatorial TFs.
b. *Tethered sliding mechanism*. This required minimal free energy input in the formation of initial loop. Although the directional dependent movement of TFs is not possible here, the free energy barrier involved in the initial dynamic loop formation stage restricts the initial landing position of DBD2s of TFs close to the promoters.
c. *Tethered binding-unbinding mode*. This mechanism is similar to the tethered sliding mode with restrictions on the sliding dynamics. Here the searching for the promoters is achieved via repeated binding-unbinding of the tethered TFs. Directional dependent movement of TFs along DNA is not possible in this mode.
d. *Parallel searching of two DBDs of TFs*. Here two different types of DBDs of the protein-protein complex of combinatorial TFs (DBD1 and DBD2) search for their cognate sites on DNA (S1 and P respectively) independently through a combination of 1D and 3D diffusions. When these DBDs binds their cognate sites simultaneously, then the looping of the intervening DNA segment automatically occurs as a result. However, this mechanism works well only for the single TF based transcription activation such as Lac I system and it is almost improbable for the combinatorial binding of TFs in the gene regulation of eukaryotes. However, this mode can be a parallel (but slow) pathway of loop formation for the above said mechanisms.

### Nonspecific binding and dissociation of TFs

In multiprotein mediated DNA looping, there is always a possibility for two different TFs interact with S1 and P respectively and the looping is mediated via protein-protein interactions among these TFs. In both propulsion and pure tethered sliding models, we have assumed that the nonspecifically bound DBD2s of CRMs-TFs do not dissociate until reaching the promoter. Nevertheless, earlier studies suggested that this assumption is valid only for the average sliding length of TFs 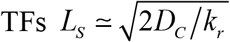, where *k*_*r*_ is the dissociation rate constant (8, 21). On can define this quantity as 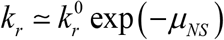 where 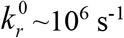 is the protein folding rate limit (65) and *μ*_*NS*_ is the free energy barrier related to the dissociation of DBD2s of TFs from DNA. Here −µ_*NS*_ is approximately the nonspecific binding energy. Clearly *μ*_*NS*_ ≥ 11 *k*_*B*_*T* is required to attain a sliding length *LS* ~ 300 bp which can be achieved via combinatorial binding of several TFs. In these calculations we assumed the 1D diffusion coefficient as *D*_*C*_ ~ 10^6^ bp^2^ s^−1^ (29, 42).

### Number of TFs required for the propulsion model

Earlier studies suggested that the DBDs of TFs behave like downhill folding proteins at their mid-point denaturation temperatures (21). Predominantly, DBDs exhibit at least two different conformational states viz. stationary and mobile (20, 21). Stationary conformational state is more sensitive to the sequence information of DNA but with less sliding mobility. Whereas, the mobile conformational state is less sensitive to the sequence information but with more sliding mobility. The average free energy barrier which separates these stationary and mobile states will be close to ~3 *k*_*B*_*T* which is approximately the energy (*μ*_*NS*_) corresponding to the nonspecific binding of single DBD with DNA i.e. the nonspecific binding energy per TF will be around −3 *k*_*B*_*T*. When there are at least four different TFs (4 DBD2s, as demonstrated in Fig. 1B) involved in a combinatorial regulation, then their cumulative nonspecific binding energy will be around −12 *k*_*B*_*T* which is sufficient to support the sliding of CRMs-TFs over ~300 bp without dissociation. In general, we have the scaling as *μ*_*NS*_ ~ 3*n* where *n* is the number of TFs involved in the combinatorial regulation. This further suggests the following relationship between *n* and *L*.

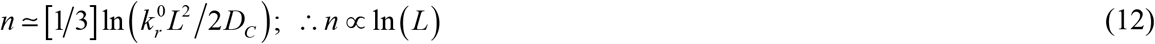

Here *L* is the distance between CRMs and promoter. Behavior of this functional relationship is demonstrated in Fig. 6. When the number TFs is less than four or distance of promoter from CRMs is higher than ~300 bp, then the CRMs-TFs complex will take either the pure tethered sliding or tethered binding-sliding-unbinding mode of dynamics to form the final synaptosome complex. Clearly, when *n* = 1 to 2 then the tethered binding-sliding-unbinding mode is the only choice for the looping mediated transcription activation.

**FIGURE 6.**
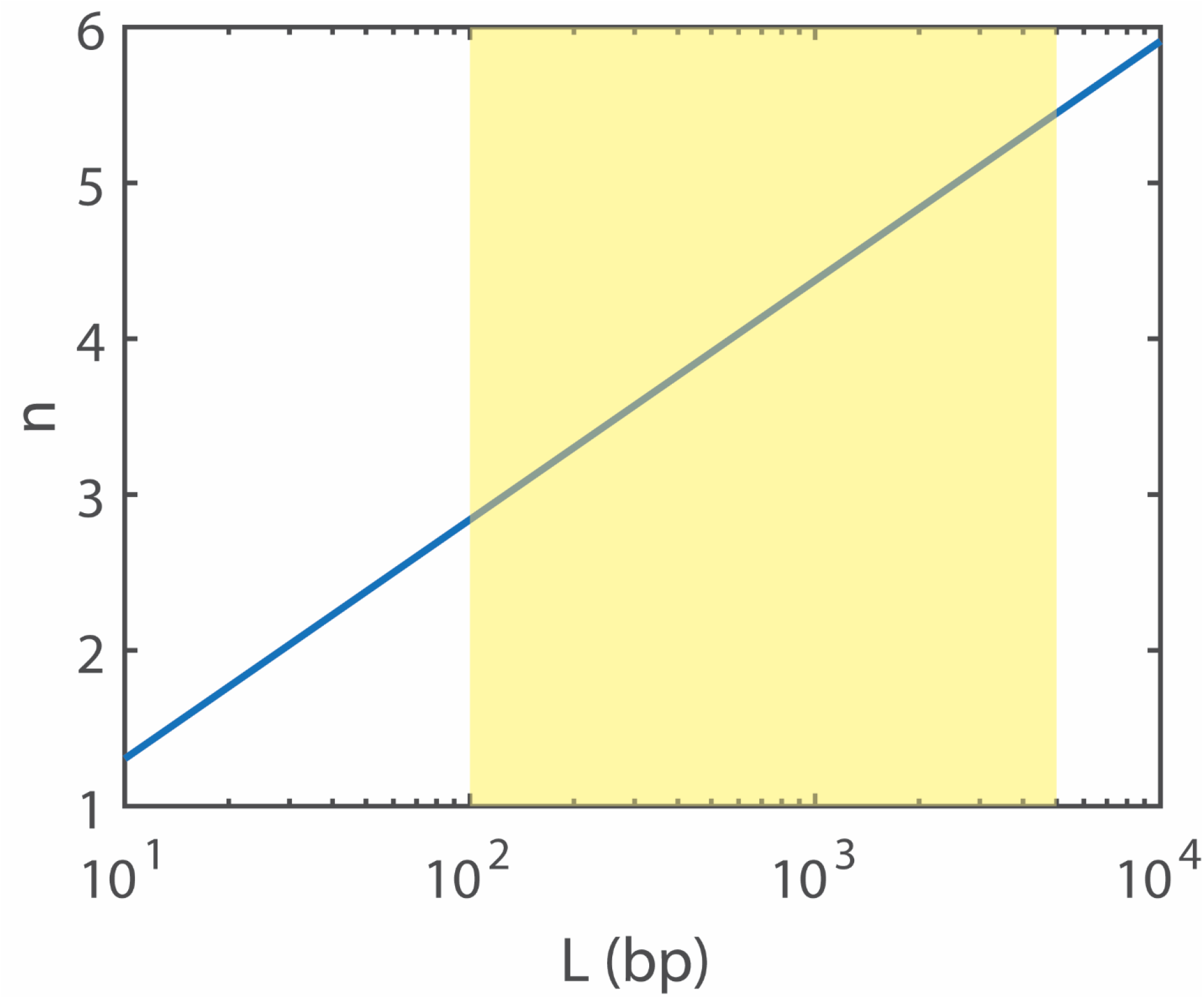
Number of combinatorial TFs required to sustain the propulsion model. Here *L* is the distance between CRMs and promoter and *n* is the number of combinatorial TFs required for the sliding of DBD2s of CRMs-TFs complex until reaching the promoter without dissociation. As in Eq. 12, 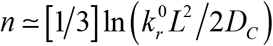. For the purpose of calculations, we have used the value for the nonspecific dissociation rate constant of DBD2s at zero free energy barrier as 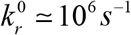 and the 1D diffusion coefficient of DBD2s of TFs on DNA as *D*_*C*_ ≃ 10^6^ bp^2^s^−1^. The yellow shaded region denotes the range of values of *L* occurring in the natural systems. These results suggest that ~3-5 TFs are enough to propel the DBD2s of TFs in most of the combinatorial regulations.

### Feasibility of the propulsion model

When the number of TFs (*n*) involved in the combinatorial regulation is ~3-5 and the average radius of gyration of individual TF is ~5 bp, then the maximum radius of gyration of the TFs complex will be around *r*_*P*_ ~15-25 bp. Our propulsion model will work only when the sequential recruitment of TFs at CRMs can bend the CRMs-DNA around them so that the radius of curvature is same as that of *r*_*P*_. This in turn requires *X*_0_ ~ 100-150 bp. When *a* ~ 150 bp, then the bending of a DNA segment of size *X*_0_ into a circular loop requires *E*_elastic_ ≃ 2π^2^*a*/*X*_0_ ~ 20-30 *k*_*B*_*T*. The average energy associated with the site-specific binding of TFs with CRMs (DBD1-S1 interactions) seems to be around –(15–20) *k*_*B*_*T* (66) and the energy associated with the cumulative nonspecific interactions of all the DBD2s of TFs (~3*n* as in Eq. 12) will be around – (10–15) *k*_*B*_*T*. When the average energy associated with the site-specific DBD1-S1 interactions via hydrogen binding network is −15 *k*_*B*_*T*, then one obtains the connection between *n* and *X*_0_ as *X*_0_ ≃ 2π^2^*a*/(15 + 3*n*). The overall specific and cumulative nonspecific binding energies of 3-5 numbers of combinatorial TFs will be around – (25–35) *k*_*B*_*T*. This amount of binding energy is more than enough not only to bend the DNA of size *X*_0_ into a circular loop around the TFs complex but also to support the sliding of DBD2s over ~300 bp without dissociation from DNA. These calculations along with Eq. 12 evidently substantiate the feasibility of the directional-dependent propulsion model.

### Limitations and Justifications

Recently, Agrawal et. al. in Ref. (56) had shown (using a two-dimensional model and numerical simulations) that the equilibrium probability associated with the protein mediated looping of a linear DNA fragment attained a minimum around *X* ~ 0.7 *a* for an end-to-end distance of ~50 bp. This minimum point seems to increase linearly with respect to the end-to-end distance of the DNA polymer. On the other hand, the equilibrium looping probability attained a maximum at *X* ~ 5*a* which was approximately independent of the end-to-end distance (56). Since the bending energy profile described by Eq. 1 decides the equilibrium probability density function associated with the loop formation, one can conclude that apart from the existence of the minimum in the bending energy profile at *X* = *X*_*C*_, there should also be a maximum in *E*_bend_ at some *X* = *X*_*D*_ as *X* tends towards zero. Therefore Eq. 1 will be valid only when the length of the DNA polymer is such that *X* > *X*_*D*_. Here one should note *X*_*C*_ ~ 13*a* which corresponds to a DNA segment with end-to-end distance equal to the length of the DNA.

In our propulsion and tethered sliding models, we have simplified our calculations by assuming circular shape for the dynamic loop structure of DNA (Figs. 1 and 3). When the loop length is comparable with that of the persistence length i.e. *a* ~ 150 bp, then the DNA segment behaves like a stiff rod. Bending such an elastic rod by pulling their terminals generally results in a tear drop or half-lemniscate like loop rather than a circular one (67). Here one should note that the bending energy at a given point on the DNA polymer is inversely proportional to the radius of curvature of that location. In a circular loop, the radius of curvature is equal in all the points. However, in the tear-drop loop, the bending energy will be higher at the focus (where the radius of curvature is less) than the terminal regions of the DNA segment. In the propulsion model, initial shape of the DNA loop well within the CRMs-TFs complex is a circular one. This possess the maximum elastic stress to propel the DBD2s of the TFs complex. As the CRMs-TFs complex slides towards the promoter, most of the elastic stresses were already released out and the lemniscate type bulging of the DNA loop will not contribute much to the propulsion of DBD2s. Therefore, it will not change our main results much. In the tethered sliding models, the dynamic loop lengths of DNA are much higher than the persistence length. Under these conditions, DNA loops take random conformations owing to increase in the chain entropy term. Therefore, assuming a circular loop of DNA is a very good approximation for the tethered sliding models.

In our calculations, we have also assumed that the TFs complex is a rigid sphere on which DNA is wrapped around. The conformational changes which occurs on the surface of the combinatorial TFs are mostly related to the cooperative protein-protein interactions associated with the recruitment processes. Since the propulsion and the tethered sliding models assume a complete assembly of all the TFs at CRMs as the initial condition, one can ignore the effects of the conformational fluctuations related to the surface level protein-protein interactions and we can consider the TFs complex as a rigid sphere.

In general, biological systems can overcome the looping energy barrier via three possible ways viz. **a**) multiprotein binding (38) which could be the origin of the combinatorial TFs in the process of evolution, **b**) placing sequence mediated kinetic traps corresponding to DBD2 in between CRMs and promoters (18) and, **c**) placing nucleosomes all over the genomic DNA to decrease the *E*_entropy_ component. All these aspects are observed in the natural systems. In multiprotein binding, the free energies associated with the DNA-protein and protein-protein interactions among TFs will be utilized in a cooperative manner for the looping of DNA. Here DBD1 and DBD2 may come from different proteins. Vilar and Saiz (38) had shown that the looping of DNA would be possible even with small concentrations of TFs when the number TFs in a combination is sufficiently large. Multiprotein binding eventually increases *X*_0_ values. However, increasing *X*_0_ will eventually decreases both the maximum possible acceleration of TFs search dynamics and the energy barrier associated with the DNA-looping. As a result, natural systems have optimized *X*_0_ between these two-opposing factors for maximum efficiency via manipulating the number of TFs involved in the combinatorial regulation.

## CONCLUSIONS

We have shown that DNA-loops can propel the complex of combinatorial transcription factors along DNA from their specific binding sites towards the promoters. The source of propulsion is the elastic energy stored on the bent DNA of the site-specifically bound DNA-protein complex. We considered directional-dependent propulsion, tethered sliding and tethered binding-sliding-unbinding models on the looping mediated transcription activation.

In directional-dependent propulsion model, the first arrived transcription factor at CRMs further recruits other TFs via protein-protein interactions. Such TFs complex has two different types of DNA binding domains viz. DBD1 which forms tight site-specific complex with CRMs of DNA via hydrogen bonding network and promoter specific DBD2s which form nonspecific electrostatic interactions around CRMs. When the overall energy associated with the specific as well as cumulative nonspecific interactions are sufficient enough, then the DNA segment around CRMs will be bent into a circle around the TFs complex. The number of TFs involved in the combinatorial regulation plays critical role here. Since the site-specific interactions at CRMs and the cumulative nonspecific interactions between DBD2s of TFs and DNA are strong enough to resist the dissociation, sliding of DBD2s well within the Onsager radius associated with the DBD2s-DNA interface towards the promoter is the only possible way to release the elastic stress of bent DNA. When the DBD2s reach the promoter without dissociation, then they form tight synaptosome complex which is necessary for the transcription activation.

When the number of TFs involved in the combinatorial regulation is not enough to bend the DNA in to a circle around the TFs complex, then one can consider tethered sliding or tethered binding-sliding-unbinding models. In the tethered sliding model, the specifically bound TFs complex at CRMs forms nonspecific contacts with DNA via dynamic loop structure and then slide along DNA without dissociation until finding the promoter. In case of tethered binding-sliding-unbinding model, the site-specifically bound TFs complex performs several cycles of nonspecific binding-sliding-unbinding before finding the promoter.

Actually, elastic and entropic energy barriers associated with the looping of DNA seem to shape up the distribution of distances between TF binding sites and promoters in the process of genome evolution. We argued that the commonly observed multiprotein binding in gene regulation might have been acquired over evolution to overcome the looping energy barrier. Presence of nucleosomes on the genomic DNA of eukaryotes might have been evolved to reduce the entropy barrier associated with the looping.

## APPENDIX A

The energy component *E*_entropy_ that is required to compensate the chain entropy loss for a Gaussian chain can be computed as follows. Let us assume that the looping of DNA occurs when 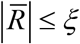 where 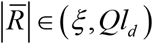(in m) is the end-to-end distance vector, ζ is the minimum looping-distance (in m) and *Ql*_*d*_ is the maximum length of the DNA polymer. Here *Q* is the total number of nucleotide monomers present in the DNA polymer and *ld* ~ 3.4 × 10^−10^ m = 1 bp is the distance between two consecutive nucleotides. The probability density function associated with the end-to-end vector 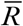 can be approximately written as 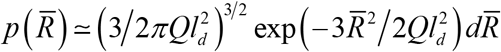(68, 69). The entropy loss upon looping of DNA is Δ*S*_*loop*_ ≃ ln(*P*_1_/*P*_Ω_ (in *k*_*B*_ units) where 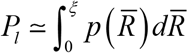 is the probability of finding the loops is and 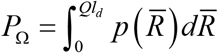 is the cumulative probability of finding all the possible configurations including loops. Explicitly one can write this down as,

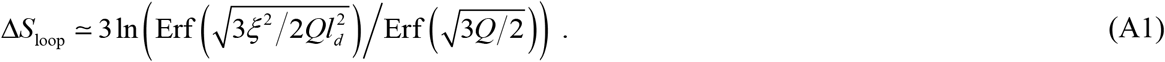

Here 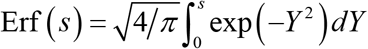 is the error function (57). When ζ ≃ *l*_*d*_ is very small then 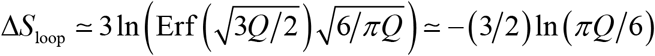 for large *Q* (52). This expression for the entropy is closely linked with the Jacobson-Stockmayer factor, or *J*-factor associated with polymer looping (37). Since *Q* = *X*/*b* where *X* is the length of the DNA polymer in bp and *b* = 1 bp is the distance between two consecutive nucleotide monomers of the DNA polymer, one finally obtains *E*_bend_ ≃ 2π^2^*a*/*X*+ (3/2)ln(π*X*/6*b*.

## Supporting Material

### 1. Theory of site-specific binding of TFs with CRMs

Let us consider a linear DNA of size *N* bps containing a *cis*-regulatory module (**CRM**) of a transcription factor (**TF**) of interest at an arbitrary location. The overall random search time or mean first passage time (**MFPT**) *τ*_*U*_ required by this TF to find its CRM via a combination of 3D and 1D diffusions can be written as follows (1, 2).

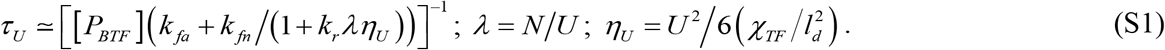

Here, [*P*_*BTF*_] (M, mols/lit) is the concentration of TF in cytoplasm, *χ_TF_* (m^2^ s^−1^) is the 1D diffusion coefficient of TFs on DNA, *k*_*fa*_ (M-^1^ s-^1^) is the Smolochowski type bimolecular rate constant associated with the direct site-specific binding of TF with CRM via 3D diffusion, *k*_*fn*_ ≃ *k*_*fa*_ (*N/R_P_*) is the overall non-specific binding rate via 3D diffusion. In this, *R*_*P*_ is the radius of gyration of TF and *k*_*r*_ (s^−1^) is the rate of dissociation of nonspecifically bound TF from DNA. Detailed theoretical studies suggested (1) an expression for *k*_*fa*_ as *k*_*fa*_ ≃ *k*_*t*_(*p*_R_δ/8). In this expression, *k*_*t*_ ≃ 8*k_B_T*/3φ (M^−1^ s^−1^) is the maximum possible 3D diffusion-limited bimolecular collision rate where *k*_*B*_ is the Boltzmann constant, *T* is the absolute temperature and *φ* is the viscosity of the aqueous medium surrounding TFs. The numerical factor 1/8 accounts for the geometry of the random coiled and relaxed conformational state of DNA. Here (1 bps = *ld* = 3.4 × 10^−10^ m). Under *in vitro* laboratory conditions (*T* = 298K and *φ* ~ 10^−1^ kg m^−1^ s^−1^ for aqueous buffer solution), one finds that *k*_*t*_ ~ 10^8^ M^−1^s^−1^ (3). Further, *p*_*R*_ is the equilibrium probability of observing a specific or nonspecific binding site to be free from other dynamic roadblocks (1, 4) which are all present on the same DNA and 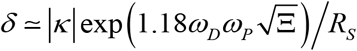 is the factor which accounts for the overall electrostatic attractive forces and the counteracting shielding effects of the solvent and other ions operating at the DNA-protein interface (1). Here *R*_*S*_ ≈ *R*_*P*_ + *R*_*D*_ is the corresponding reaction radius, *R*_*D*_ is the radius of the DNA cylinder and |κ| is the ***Onsager radius** which is defined as the distance between charged reactant molecules at which the overall electrostatic energy will be the same as that of the background thermal energy*. Further ω_*D*_ and ω_*P*_ are the overall charges on the DNA backbone and the DNA binding domains (**DBDs**) of TFs respectively and Ξ is the ionic strength of the surrounding reaction medium.

The term *λ* in **Eq. S1** is the minimum number of association-scan-dissociation cycles required by TFs to scan the entire DNA sequence of size *N* (bps) and η_U_ is the (averaged over initial landing positions) overall MFPT that is required to scan *U* bps of DNA via pure 1D diffusion. Here *U* (bps) is a random variable which takes different values in each of the association-scan-dissociation cycles. The probability density function associated with the 1D diffusion lengths *U* of TFs can be written as follows (1).

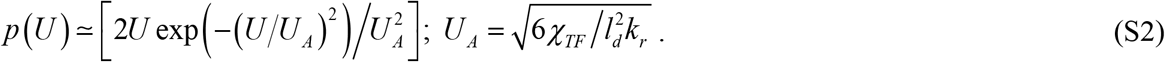

Here *U*_*A*_ is the maximum achievable 1D diffusion length of the nonspecifically bound TFs on DNA that is measured in bp. When TF moves with a jump size of *k* (bps) then we find that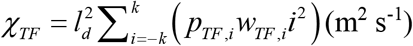 where *i* ∈ *Z*, *w_TF_, ±i* are the microscopic transition rates (1/*s*) associated with the forward and reverse movements of TFs with jump size of *i*, and *p_TF_, ±i* are the corresponding microscopic transition probabilities (5, 6). Here the dynamics of TF with jump size *k* means that from the current location *x* (measured in m) TF can hop anywhere inside (*x* – *k l*_*d*_, *x* + *k l*_*d*_) with equal probabilities i.e. 1/2*k*. The step length *k* is measured in bp. Since the dynamics at the DNA-protein interface involves segmental motion of DBDs of TFs, one can assume the protein folding rate limit (7) for *w*_*TF*_, *±1* ~ 10^6^ s^−1^. Noting that *p_TF_, ±1* ~ ½ for an unbiased 1D random walk, one finds that *χ_TF_*/*l*_*d*_^2^ ~ 10^6^ bps^2^s^−1^ corresponding to the sliding of TFs on DNA for which *k* = 1 bps. Approximately this is the experimental value of the 1D diffusion coefficient associated with the sliding of TFs on DNA (1, 8, 9). Using the probability density function corresponding to the sliding length, one can define the overall average search time as 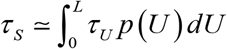.

For an arbitrary jump size *k* with the average microscopic transition probabilities as *p*_*TF*,±*i*_ = 1/2*k* and average transition rates 〈*w*_*TF,i*_〉 = φ_*TF*_, one finds that 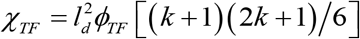. Recently, Amitai has shown (10) that *p*_*TF,±i*_ ∝*i*^−1/2^ and *w*_*TF*_, _±*i*_ ∝ *p*_*TF,±i*_ (Eqs. 2, 4-5 of Ref. (10) where we have |*n*-*l*| = *i* in the present context) so that (*p*_*TF,±i*_ *W*_*TF*,±*i*_)*∝i*^−1^ for large size chromatin DNA. Upon implementing this scaling, one obtains that χ_*TF*_ ∝ [*k*(*k* +1)/2]. Therefore, our formula for χ_TF_ with respect to *k* is asymptotically consistent with Ref. (10). Here the jump size *k* is positively correlated with the degree of condensation of DNA. Densely packed DNA allows larger jump sizes. The 3D diffusion coefficient of TFs, i.e. *χ*^3D^ will be always ~10-10^2^ times higher than *χ^TF^* and in general *χ_TF_* ≤ *χ_3D_* irrespective of *k* and presence of various 1D facilitating processes such as hopping and intersegmental transfers. Various symbols and parameters used throughout this supporting material as well as in the main text are listed in Table S1.

Upon mixing DNA with TF in buffer, the site-specific complex can be formed via two different pathways viz. (**a**) direct site-specific binding through 3D diffusion and (**b**) through a combination of 1D and 3D diffusions. One can describe this by a pseudo first order scheme 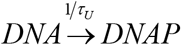. Here the concentration of specific site will be equal to the concentration of DNA, and DNAP is the site-specific complex. In *τ_U_* of **Eq. S1**, [*P*_*BTF*_]*k*_*fa*_ is the pseudo first order rate of direct site-specific binding of TF with DNA via pure 3D diffusion and *k*_*fn*_[*P*_*BTF*_]/(1 + *k*_*r*_*λη_U_*) is the pseudo first order rate of site-specific binding of TF via a combination of 1D and 3D diffusion.

### 2. Looping mediated communication between CRMs-TFs with the promoter

The mean first passage time (MFPT) associated with the DNA binding domains 2 (DBD2) of CRMs-TFs complex to reach the promoter of a gene via *DNA-loop mediated propulsion mechanism* while the DBD1 of TF complex is still tightly bound with S1 of DNA (Fig.1 of the main text and Fig. S1 for details) can be given as follows.

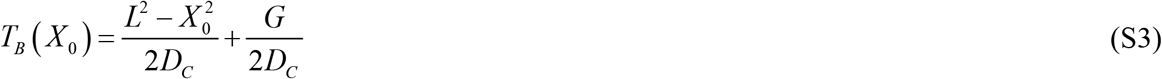

Here *X_0_* is the initial position of DBD2s of CRMs-TFs on DNA or the initial loop length, *L* is the location of promoter, *a* is the persistence length of DNA and *D*_*C*_ is the one-dimensional diffusion coefficient associated with the sliding of DBD2s of TFs along DNA. We have assumed here that the site-specific DBD1-S1 is strong and intact (Fig. 1B of the main text). The function *G* can be defined as follows.

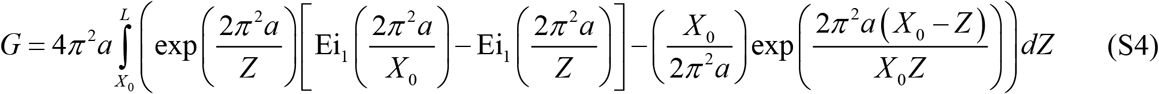

Here 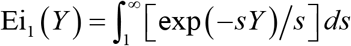 is the E_1_ exponential integral (11).

Noting that *T*_*N*_(*X*_0_) = (*L* − *X*_0_)^2^/2*D*_*C*_ (from Eq. 7 of the main text) which is the MFPT associated with the finding of promoter by TFs complex starting from *X_0_* via pure sliding dynamics without dynamics DNA loop structures, one can define *η_P_* which is the number of times the dynamics loop driven sliding of CRMs-TFs towards the promoter is faster than the normal 1D sliding dynamics as follows.

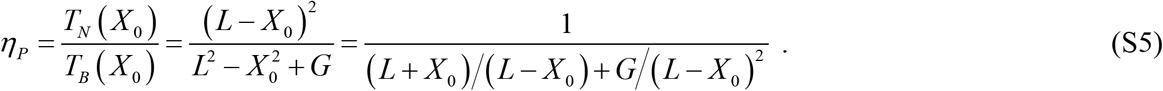

Clearly *η_P_* is not dependent on *D*_*C*_ and it depends only on the parameters (*L*, *X_0_* and *a*). Further, 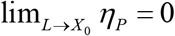 since *T*_*N*_(*X*_0_) approaches zero much faster than *T*_*B*_(*X*_0_) as *L* tends towards infinity (see Fig. S2 for details). There also exists an asymptotic limit as lim_*L*→∞_ η_P_ = 1. This means that 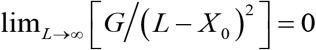 (see Fig. S3 for details). The optimum distance between CRMs and promoter i.e. *L*_opt_ at which *η_P_* is a maximum can be obtained by solving *dη_P_*/*dL* = 0 for *L* for given *a* and *X_0_*. Explicitly one can write down this as follows.

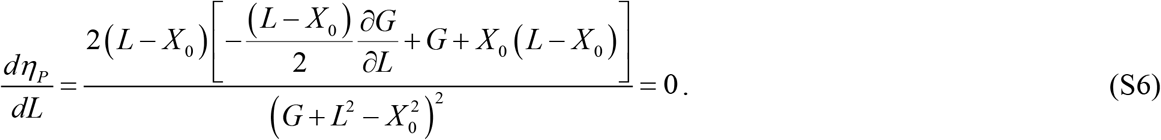

This has a trivial solution *L* = *X_0_*. Upon ignoring this one, *L*_opt_ can be obtained by numerically solving the following equation for *L* at given *a* and *X_0_*.

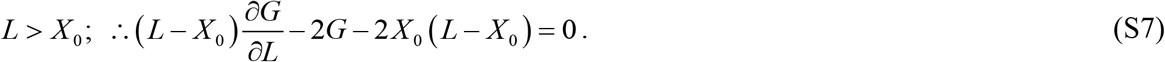

---

**FIG S1.**
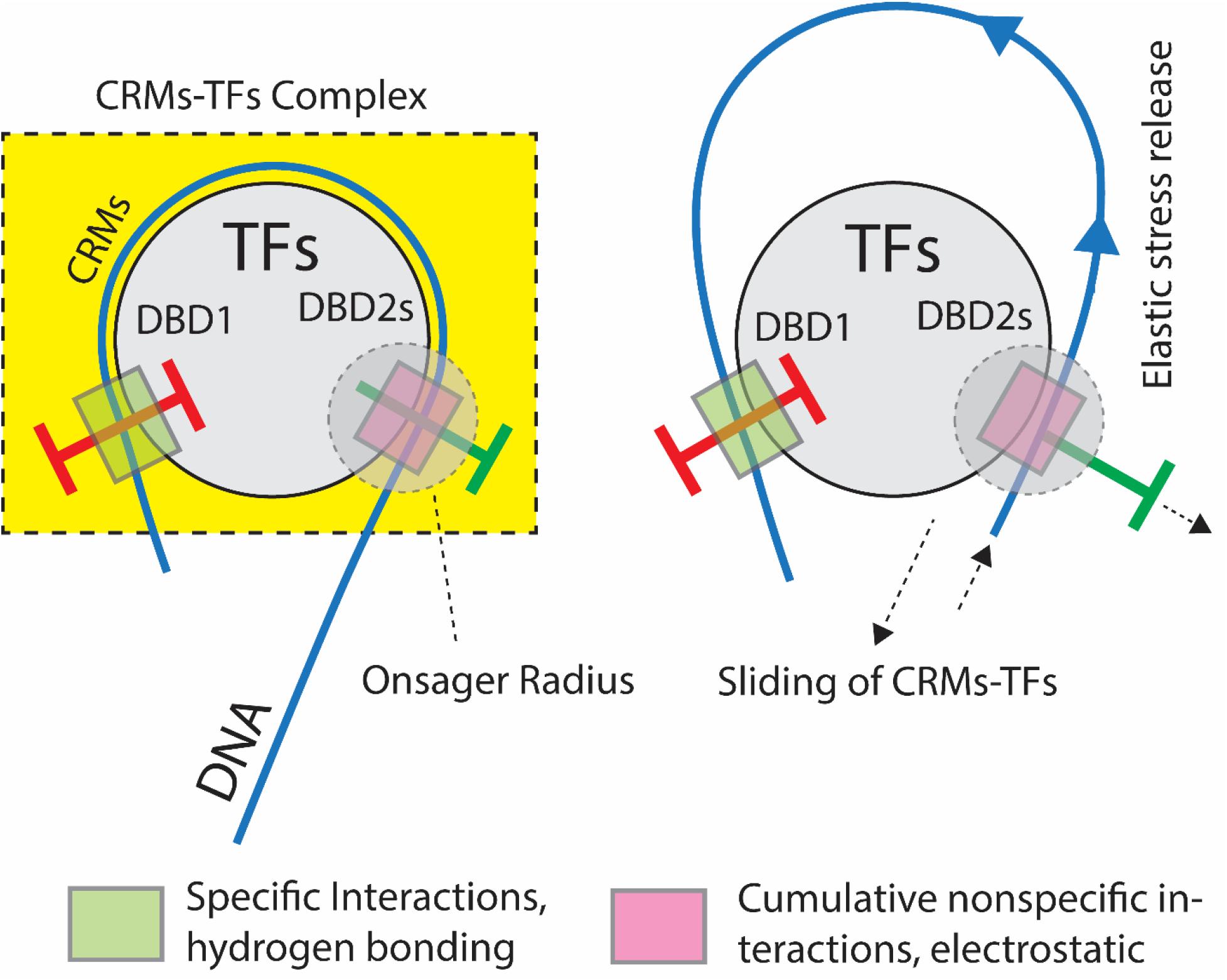
Schematic diagram on the mechanism of propulsion of CRMs-TFs on DNA. In the propulsion model, Combinatorial TFs assemble on the CRMs via recruitment mechanism and bend the DNA around final TFs complex. This TFs complex has two different DBDs viz. DBD1 which site-specifically interacts with the CRMs via hydrogen bonding network and several DBD2s which interact with the DNA near CRMs via nonspecific electrostatic interactions. Both DBD1-CRMs as well as DBD2s-DNA interactions are strong enough to keep the bent DNA intact. In this scenario, the elastic stress of bent DNA of CRMs-TFs complex can be released only via sliding of DBD2s of TFs along DNA well within the Onsager radius associated with the DBD2s-DNA interface. Propulsion mechanism works only when the cumulative nonspecific interactions are strong enough to keep the DBD2-DNA well within the Onsager radius until CRMs-TFs reaches the promoter. This in turn requires the binding of several combinatorial TFs.

**FIG S2.**
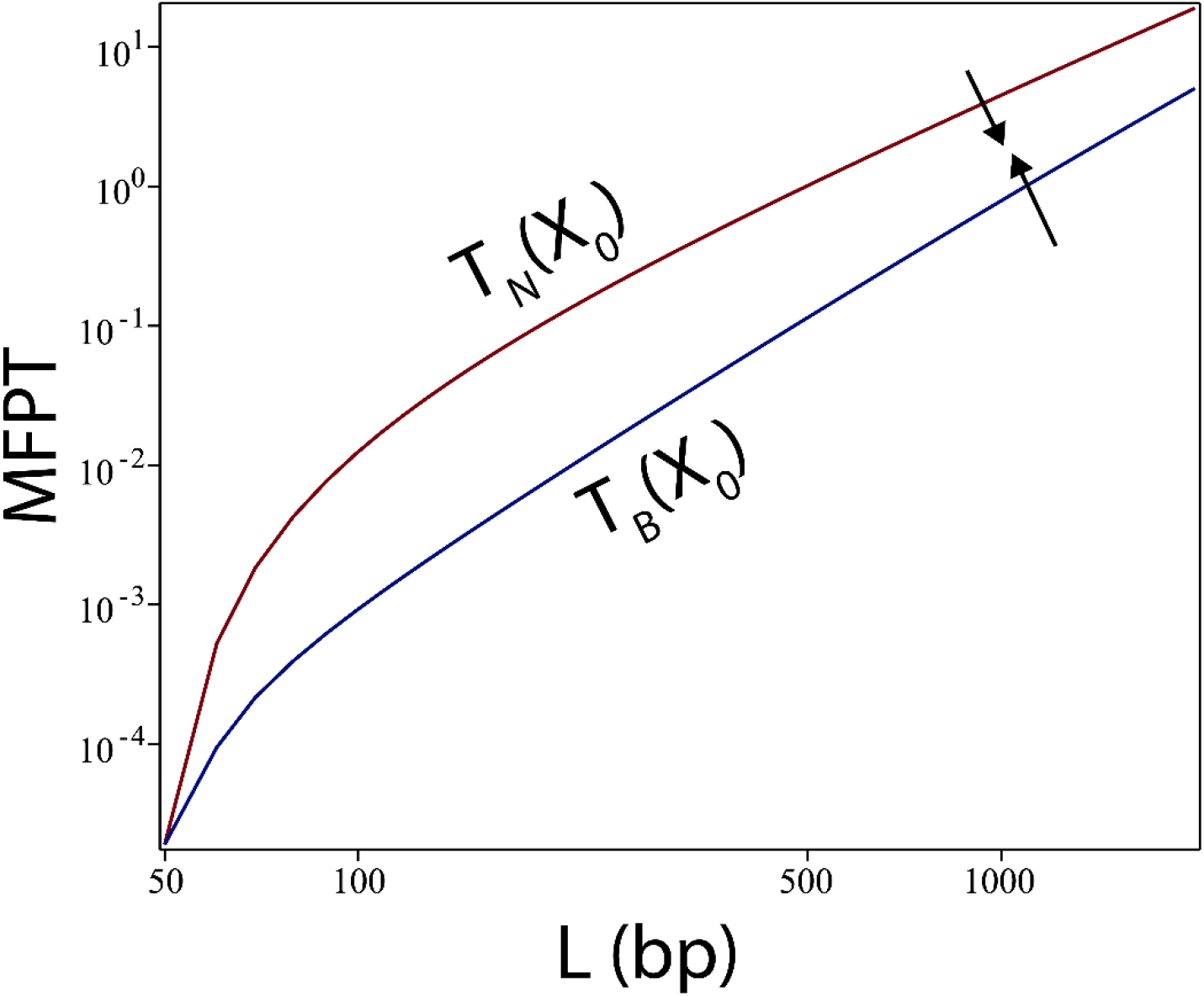
Showing that *T*_*N*_(*X*_0_) approaches zero much faster than *T*_*B*_(*X*_0_) so that 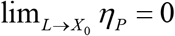. Here the settings are *X*_0_ ~ 50 bp and *a* ~ 150 bp and *L* was iterated from 50 to 2000 bp. Further we also find the limit lim_*L→∞*_*T*_*B*_(*X*_0_) = *T*_*N*_(*X*_0_) or 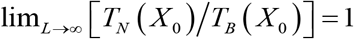.

**FIG S3.**
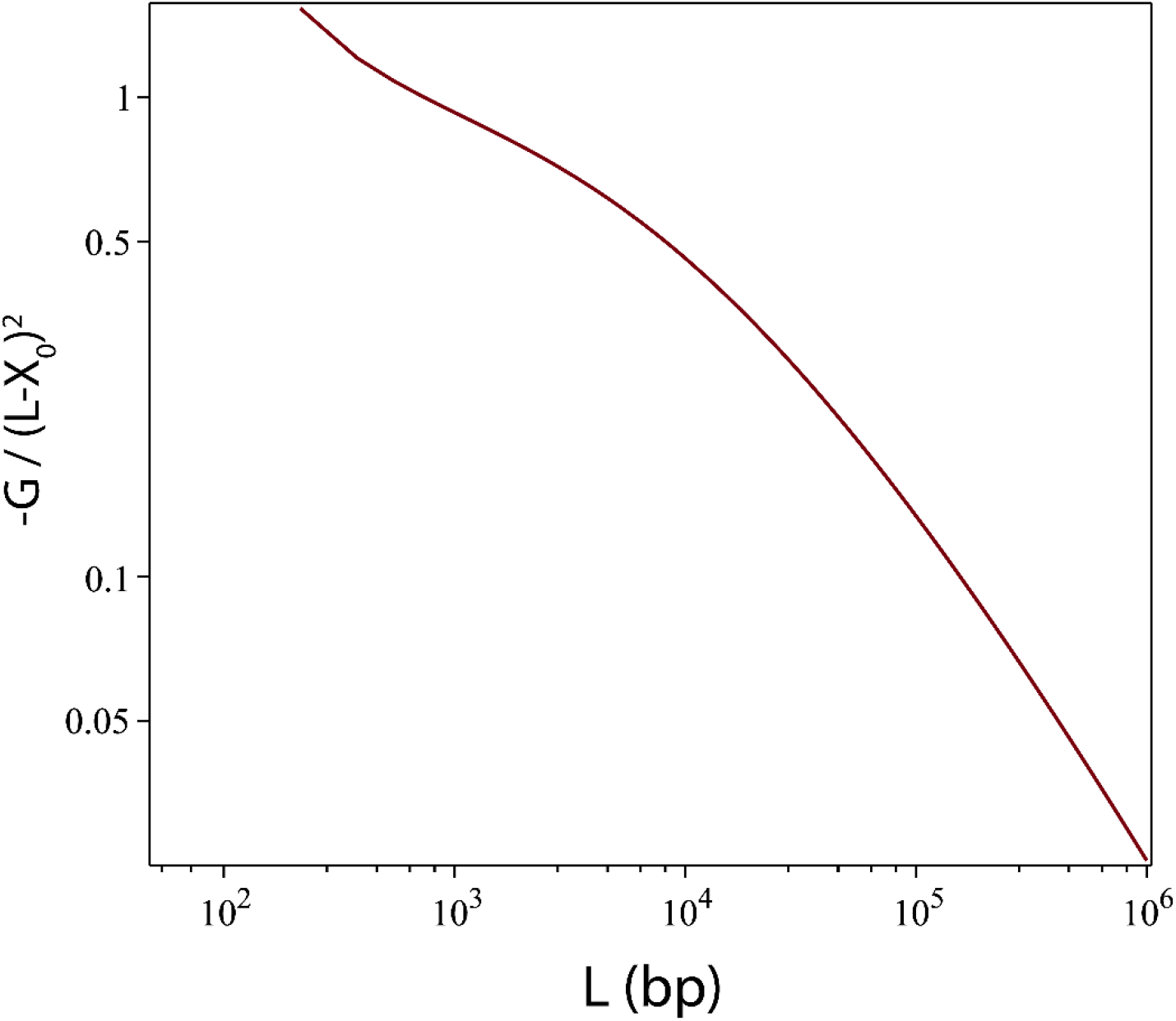
Showing 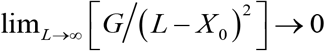. Here the settings are *X*_0_ ~ 50 bp and *a* ~ 150 bp and *L* was iterated from 50 to 10^6^. The function *G* is defined as in **Eq. S4**.

**FIG S4.**
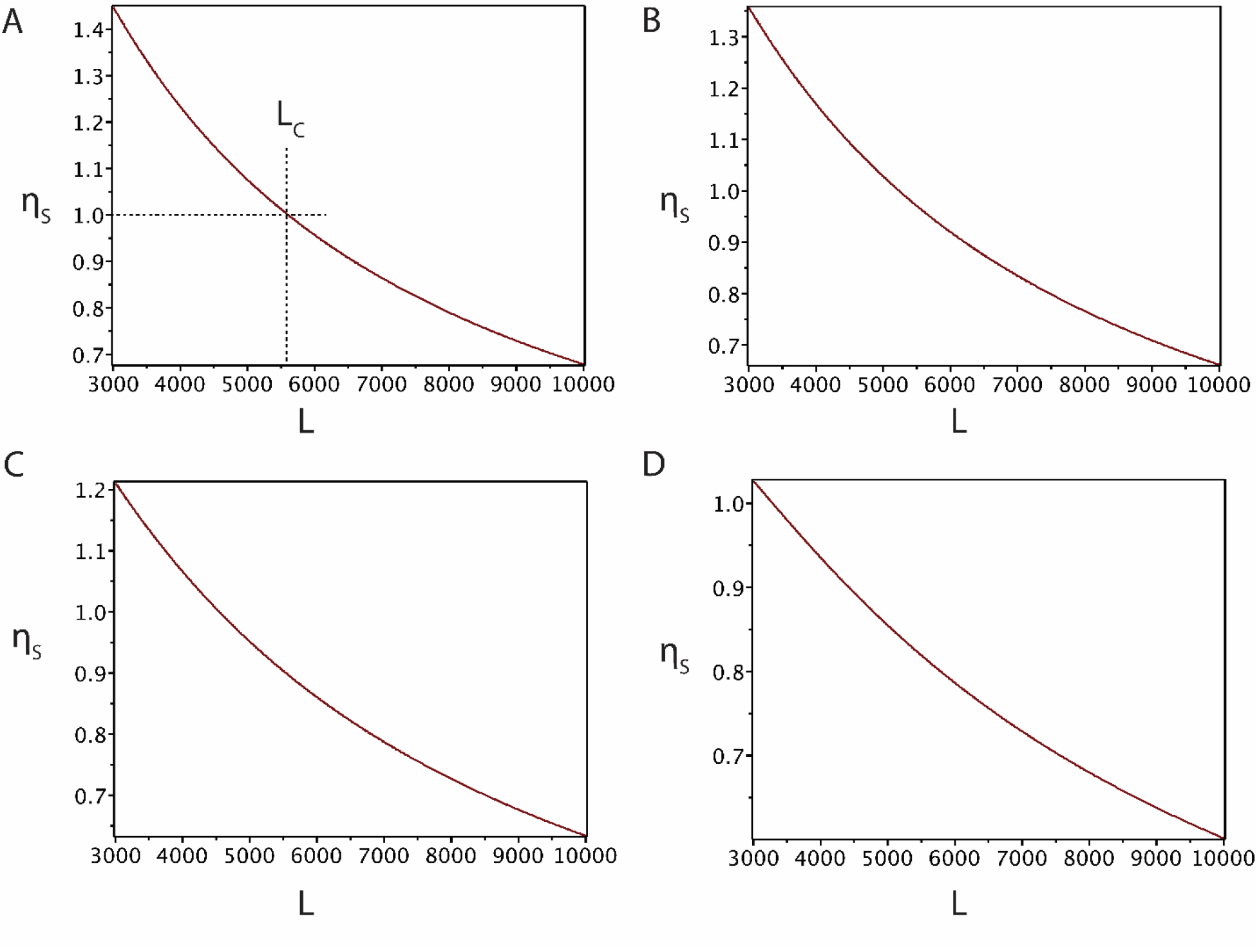
Variation of the critical distance (*L*_*C*_) between CRMs and promoter in the tethered sliding model. Here η_S_ = *T*_*N*_ (*X*)/*T*_*U*_(*X*) as defined in Eqs. 7 and 9. In these MFPTs, *X* is the initial loop length, *X*_0_ is the left side reflecting boundary and *L* is the right-side absorbing boundary. *L*_*C*_ is defined such that when *L* < *L*_*C*_ then *η*_*S*_ > 1 and when *L* > *L*_*C*_ then *η*_*S*_ < 1. Settings are (see Eq. 9 of the main text), *X* = *XC* where *X*_*C*_ = 4π^2^*a*/3, persistence length of DNA *a* ~ 150 bp and *L* was iterated from 3000 to 10000 bp. **A**. *X*_0_ = 100 bp. **B**. *X*_0_ = 250 bp. **C**. *X*_0_ = 500 bp. **D**. *X*_0_ = 1000 bp. These results suggest that *L*_*C*_ decreases as *X*_0_ increases. For the most probable value of *X*_0_ ~ 100 bp one finds that *L*_*C*_ ~ 3*X*_*C*_.

**FIG S5.**
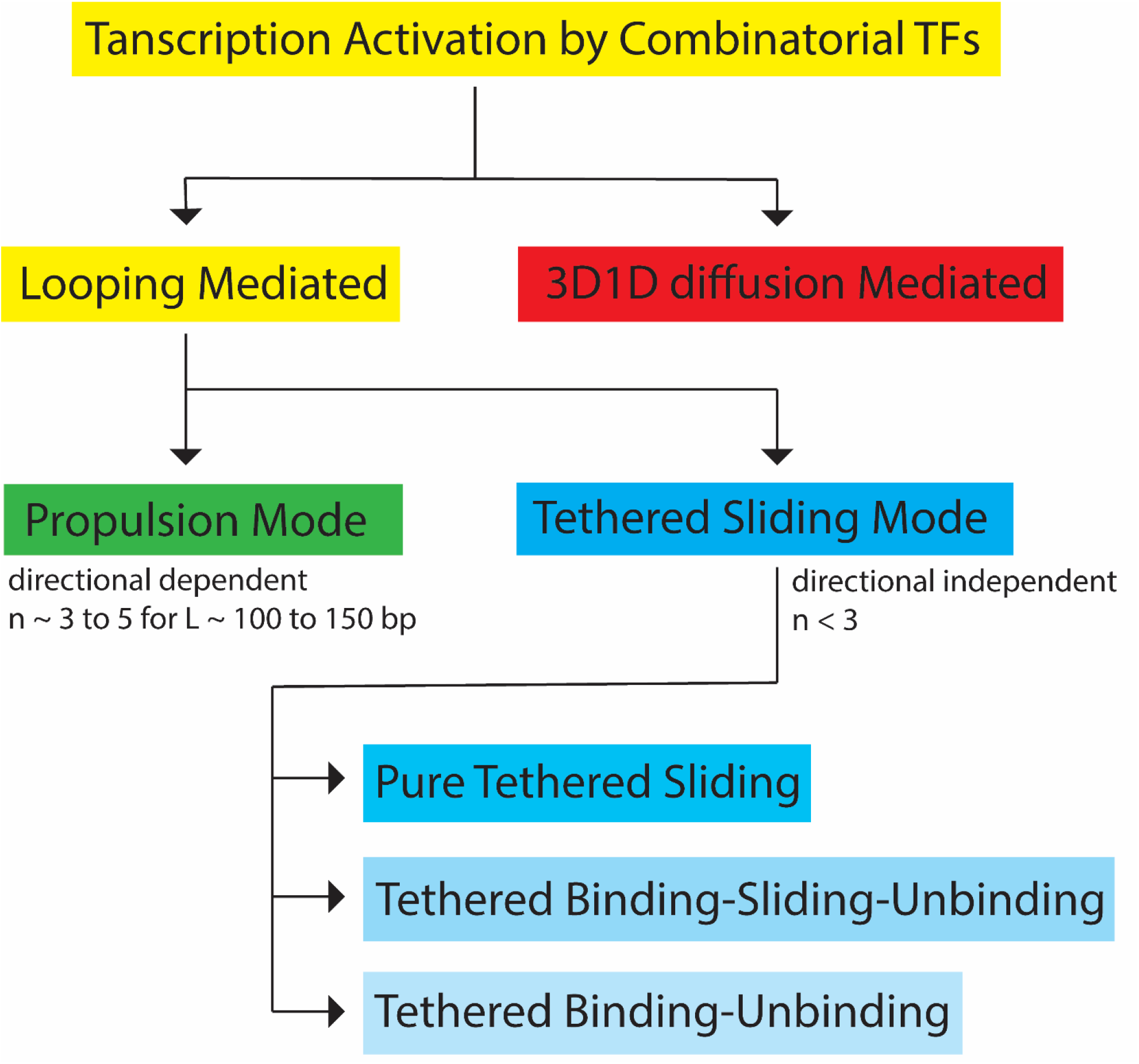
Summary of various models proposed on the transcription activation of combinatorial TFs. Looping mediated activation can be via the directional dependent propulsion mode or tethered sliding mode depending on the number of combinatorial TFs. Here *n* is the number TFs involved in the combinatorial regulation and *L* is the distance between CRM and promoter. Propulsion model will work only when the overall energy associated with the specific and nonspecific binding of combinatorial TFs is enough to bend the CRMs-DNA around the TFs complex and also sustain the continuous sliding of CRMs-TFs complex until reaching the promoter region. Tethered binding-unbinding model has been studied in detail in Ref. (12).

**TABLE S1.**
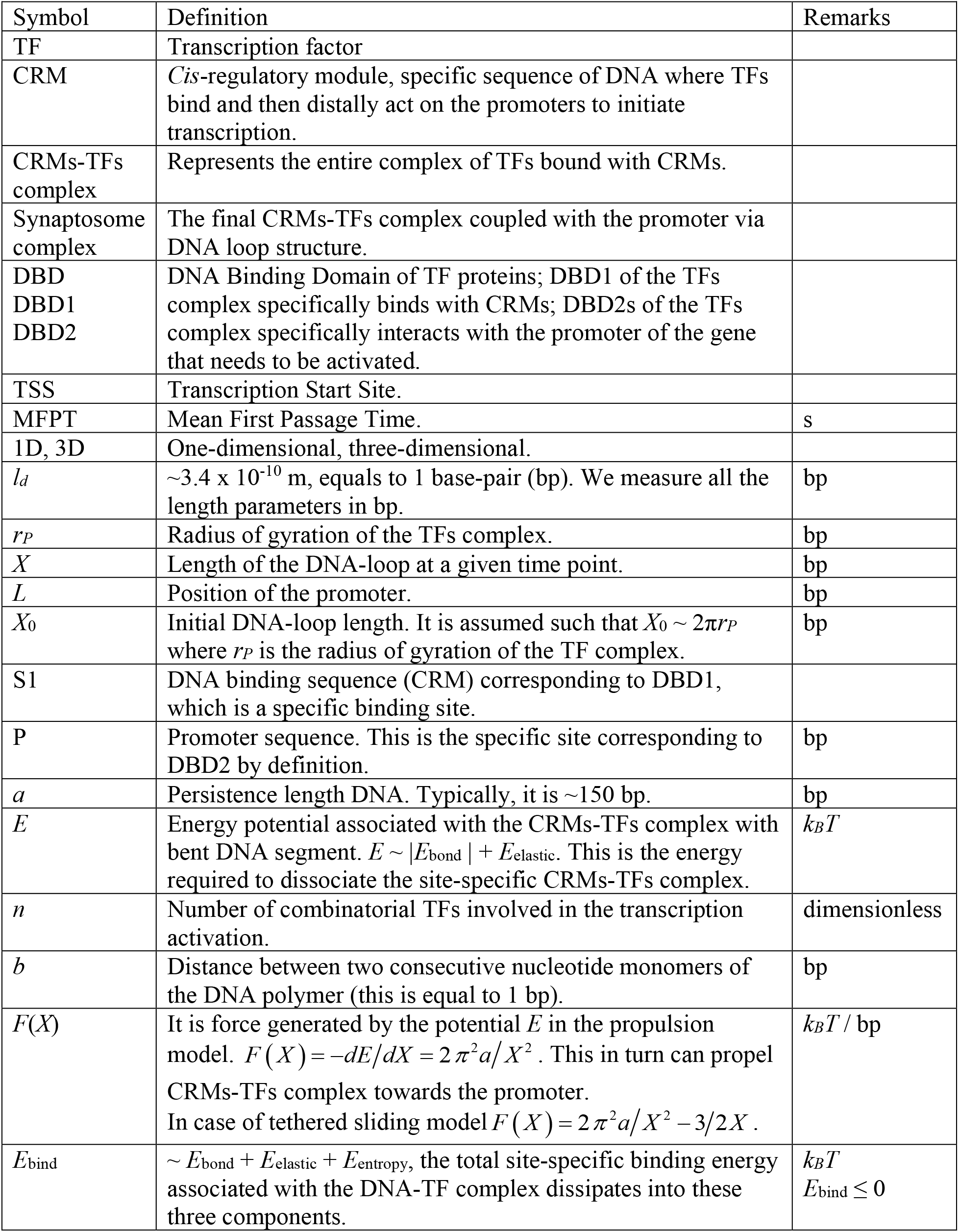
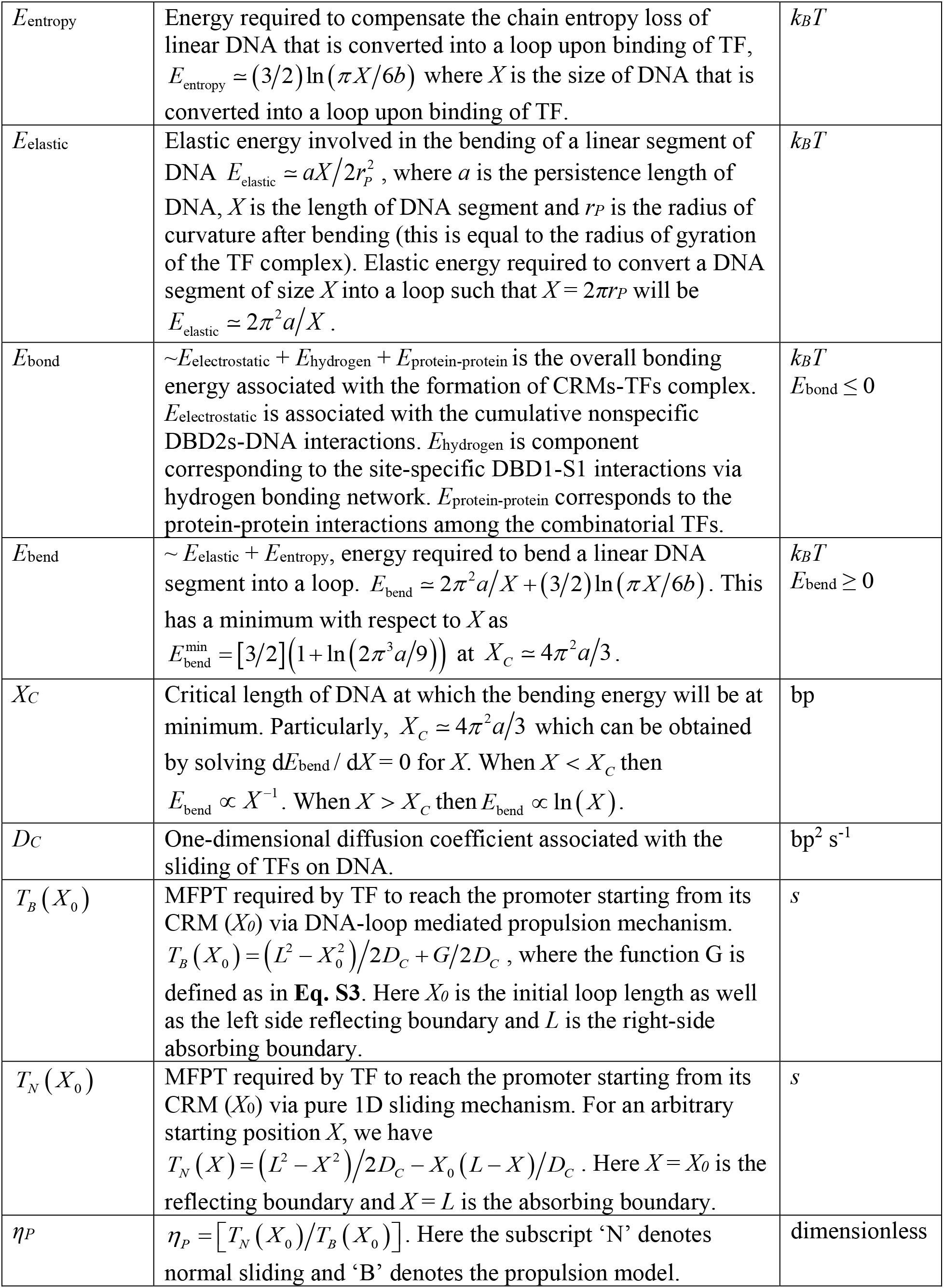
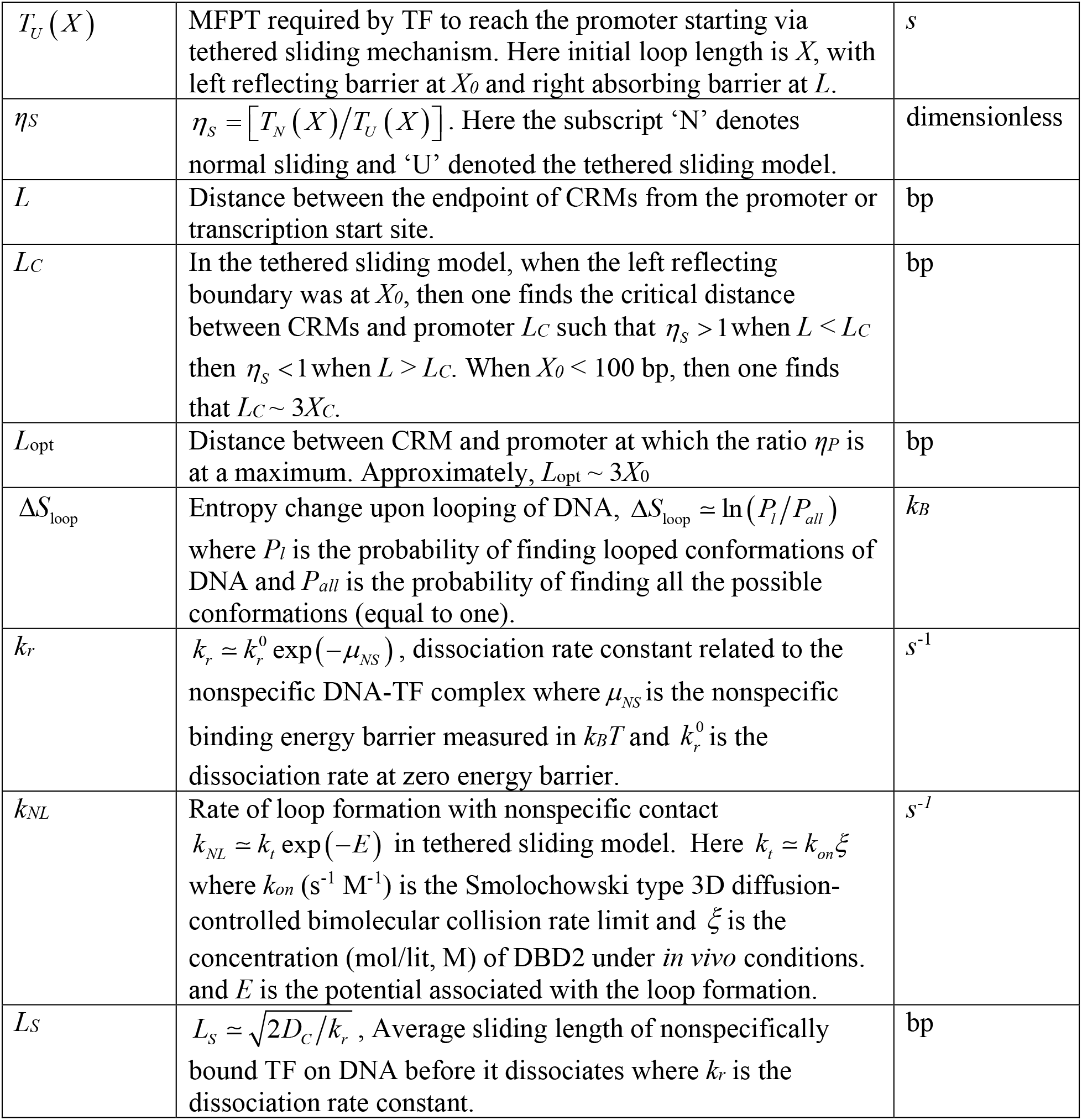
List of symbols used in the main text

